# RELAX-Jr: An Automated Pre-Processing Pipeline for Developmental EEG Recordings

**DOI:** 10.1101/2024.04.02.587846

**Authors:** Aron T. Hill, Peter G. Enticott, Paul B. Fitzgerald, Neil W. Bailey

**Affiliations:** Cognitive Neuroscience Unit, School of Psychology, Deakin University, Geelong, Australia; Monarch Research Institute, Monarch Mental Health Group, Sydney, Australia; School of Medicine and Psychology, The Australian National University, Canberra, ACT

**Author notes:** Correspondence: Aron T. Hill, PhD Cognitive Neuroscience Unit School of Psychology Deakin University.

**Keywords:** EEG, development, child, preprocessing, pipeline, automated, artifact, cleaning

## Abstract

Automated EEG pre-processing pipelines provide several key advantages over traditional manual data cleaning approaches; primarily, they are less time-intensive and remove potential experimenter error/bias. Automated pipelines also require fewer technical expertise as they remove the need for manual artifact identification. We recently developed the fully automated Reduction of Electroencephalographic Artifacts (RELAX) pipeline and demonstrated its performance in cleaning EEG data recorded from adult populations. Here, we introduce the RELAX-Jr pipeline, which was adapted from RELAX to be designed specifically for pre- processing of data collected from children. RELAX-Jr implements multi-channel Wiener filtering (MWF) and/or wavelet-enhanced independent component analysis (wICA) combined with the adjusted-ADJUST automated independent component classification algorithm to identify and reduce all artifacts using algorithms adapted to optimally identify artifacts in EEG recordings taken from children. Using a dataset of resting-state EEG recordings (N = 136) from children spanning early-to-middle childhood (4-12 years), we assessed the cleaning performance of RELAX-Jr using a range of metrics including signal-to-error ratio, artifact-to-residue ratio, ability to reduce blink and muscle contamination, and differences in estimates of alpha power between eyes-open and eyes-closed recordings. We also compared the performance of RELAX-Jr against four publicly available automated cleaning pipelines. We demonstrate that RELAX-Jr provides strong cleaning performance across a range of metrics, supporting its use as an effective and fully automated cleaning pipeline for neurodevelopmental EEG data.

## Introduction

Almost a century on from Hans Berger’s seminal non-invasive recordings of electrical brain activity in humans (Berger, 1929), electroencephalography (EEG) remains a highly popular method for investigating functional brain dynamics in both health and disease. Key advantages of EEG are its temporal precision (millisecond range) and cost-effectiveness. These advantages have facilitated its integration into broad-ranging experimental paradigms and clinical practice, where it is frequently used to interrogate both spontaneous and task-related neural activity. The wide availability and affordability of EEG also makes it highly conducive to large-scale cross-sectional and longitudinal investigations into typical neurodevelopment, as well as for studying aberrant neural activity patterns across a range of neurodevelopmental disorders (Buzzell et al., 2023; Cellier et al., 2021; Gabard-Durnam et al., 2019; Marshall et al., 2002).

EEG data, however, are frequently contaminated by a number of artifacts, both biological and environmental, which, if not adequately removed or suppressed, can obscure the underlying neural signals of interest. Furthermore, many of the biological artifacts, such as electromyographic (EMG) activity, movement related interference, and eye blinks/ocular artifacts, are often more common and pronounced in developmental populations (Brooker et al., 2020; Herve et al., 2022). Additional common artifacts (that are not specific to children) include electrical line noise at either 50 or 60 Hz, electrocardiographic (ECG) related signals, and noise related to poor electrode impedance (Daube, 2009; Sazgar & Young, 2019; Tandle & Jog, 2016). Pre-processing strategies for reducing these artifacts can vary widely, both between investigators, and across laboratories, with a growing array of EEG artifact removal algorithms available to investigators (Jiang et al., 2019; Roy et al., 2021). Manual detection (via visual inspection) and removal of artifacts is frequently performed; however, this process is time-consuming, subjective, often imprecise, and not easily scalable to large datasets (Delorme, 2023; Mumtaz et al., 2021). Manual artifact reduction also requires considerable operator expertise in order to accurately interpret the EEG signal, making it vulnerable to potential bias and inconsistency resulting from human influence (Fitzgibbon et al., 2007; Jas et al., 2017).

Advances in signal processing methods have seen automated and semi-automated cleaning approaches become increasingly utilised in EEG pre-processing. These automated approaches have the benefit of reducing experimenter error and/or bias, improving efficiency, and creating transparent and replicable workflows, thus advancing reproducible and open science (De Blasio & Barry, 2023). To this end, we previously released the open-source Reduction of Electroencephalographic Artifacts (RELAX) software, which enabled fully automated pre-processing of EEG data. RELAX showed favourable results for cleaning resting-state and task-related datasets from adult populations when compared to several other commonly used automated pipelines (Bailey et al., 2023a; Bailey et al., 2023b). Development of RELAX was motivated by the shortage of fully automated pipelines available to the EEG community, as well as our experience that some data are not effectively cleaned (or are over-cleaned) using existing independent component analysis (ICA)- based approaches (Bailey et al., 2023a; Dimigen, 2020). In addition to RELAX, several other semi- and fully-automated pipelines for pre-processing EEG data have been developed in recent years (Chang et al., 2020; Gabard-Durnam et al., 2018). While these have been mostly targeted for use in adult populations, some recent pipelines have been developed for use with data collected in children (e.g., Debnath et al., 2020; Flo et al., 2022). However, many automated EEG pre-processing pipelines (including RELAX) have been tested only in adult data. EEG is often also used to assess neural activity across developmental populations (both typically developing and clinical), which can pose additional challenges for data cleaning. For instance, movement and muscle-related artifacts are often more apparent in children (Herve et al., 2022), while their limited attentional capabilities often necessitate shorter recording times and less cognitively demanding tasks (Bell & Cuevas, 2012; DiStefano et al., 2019; Herve et al., 2022).

Here, we present the RELAX-Jr (Reduction of Electroencephalographic Artefacts for Juvenile Recordings) EEG pre-processing software pipeline, which we have specifically developed for cleaning of EEG data recorded from children. This pipeline is based on the original RELAX software (Bailey et al., 2023a; Bailey et al., 2023b), which implements multi-channel Wiener filters (MWF) (Borowicz, 2018; Somers et al., 2018) and/or wavelet-enhanced independent component analysis (wICA) (Castellanos & Makarov, 2006) to reduce or remove artifactual signals. However, RELAX-Jr also includes important modifications, such as the inclusion of the ‘adjusted-ADJUST’ independent component analysis (ICA) artifact component selection algorithm designed for Geodesic electrode nets, which are frequently used in EEG recordings in children (Leach et al., 2020), and the use of the Preconditioned ICA for Real Data (PICARD) algorithm for maximum likelihood independent component analysis (ICA) (Ablin et al., 2018a, 2018b). Adjusted-ADJUST is more sensitive to the increased noise often present in data collected from children, and also considers a larger range of frequencies to identify alpha peaks in the neural signal, as alpha peak frequencies are often lower in children (Leach et al., 2020; Marshall et al., 2002). PICARD has been shown to perform similarly to the popular Infomax ICA algorithm (Bell & Sejnowski, 1995), but with much faster speed of convergence (Frank et al., 2022). Like RELAX, RELAX-Jr is fully automated, meaning that no user input is required after initial cleaning settings are determined, thus promoting streamlined and consistent/unbiased batch processing of data files. By default, RELAX-Jr receives as inputs raw datafiles in EEGLAB format (Delorme & Makeig, 2004), and outputs cleaned continuous data referenced to the robust average reference ready for further segmentation (optional) and analysis (Bigdely-Shamlo et al., 2015).

In this paper, we apply and rigorously test four separate versions of the RELAX-Jr pipeline, testing four specific parameter variations. These tests were implemented using a large heterogeneous test dataset of resting-state EEG (both eyes-open and eyes-closed recordings) taken from typically developing children (4- 12 years of age). We compare the results obtained from RELAX-Jr against four popular automated pre- processing pipelines: the Harvard Automated Processing Pipeline for Electroencephalography (HAPPE) (Gabard-Durnam et al., 2018), the Maryland analysis of developmental EEG (MADE) (Debnath et al., 2020), the Automated Pipeline for Infants Continuous EEG (APICE) (Flo et al., 2022), and the Artifact Subspace Removal followed by ICA (ASR) approach (Chang et al., 2020). We show that RELAX-Jr exhibits robust efficacy across a range of cleaning metrics, establishing it as an effective and unbiased automated method for processing EEG data collected from children.

## Methods

### Dataset

Each pre-processing pipeline was applied to a dataset of resting-state EEG recordings collected from a cohort of typically developing children spanning early-to-middle childhood (N = 136, age range: 4–12 years; 71 male; average age = 9.42 years, SD = 1.95). All participants were English speaking, and none had received a formal diagnosis of any neurological, psychiatric, or genetic disorder. Ethical approval was provided by the Deakin University Human Research Ethics Committee (2017–065), while approval to approach public schools was granted by the Victorian Department of Education and Training (2017_003429). The EEG data were recorded in a dimly lit room using a 64-channel HydroCel Geodesic Sensor Net (Electrical Geodesics, Inc, USA) containing Ag/AgCl electrodes surrounded by electrolyte-wetted sponges. Data were acquired either at Deakin University (Melbourne, Australia), or in a quiet room at the participant’s school using NetStation software (version 5.0) via a Net Amps 400 amplifier using a sampling rate of 1 KHz. Electrode Cz was used as the online reference. Electrode impedances were checked to ensure they were < 50 KOhms prior to recording. The data were recorded for two minutes while participants sat with their eyes open and focussed their gaze at a fixation cross on a computer screen, and two minutes while participants had their eyes closed. Three of the 136 participants did not have complete eyes-open recordings and were not included for this dataset.

### RELAX-Jr Pipelines

To optimise the RELAX-Jr pipeline, we tested adaptations of four specific versions of RELAX that were found to be most effective for artifact removal in our adult datasets: *MWF wICA*, *MWF Only*, *wICA ADJUST*, and *ICA Subtract* (Bailey et al., 2023a). We note that within the original version of RELAX (designed for adult recordings) the *wICA* and *ICA Subtract* methods used ICLabel to identify artifact components, while RELAX-Jr replaces ICLabel with adjusted-ADJUST (which is designed for use in paediatric EEG data). Key details of each of these methods are summarised in Table 1. For brevity, we have not included a detailed description of each method. For specific details, the reader is encouraged to refer to our previous publications (Bailey et al., 2023a; Bailey et al., 2023b).

**Table 1.** Summary of the steps for each of the four implemented RELAX-Jr pipelines.

|  | **MWF wICA** | **ICA Subtract** | **wICA ADJUST** | **MWF Only** |
| --- | --- | --- | --- | --- |
| **Filter** | 0.25-80 Hz Bandpass, 47-53 Hz notch filter | 0.25-80 Hz Bandpass, 47-53 Hz notch filter | 0.25-80 Hz Bandpass, 47-53 Hz notch filter | 0.25-80 Hz Bandpass, 47-53 Hz notch filter |
| **Bad Channel Rejection** | PREP's 'findNoisyChannels', then for each 1s epoch, reject if >5% of data show extreme values (defined in Fig. 1) or log-power log-frequency slopes >-0.59 | PREP's 'findNoisyChannels', then for each 1s epoch, reject if >5% of data show extreme values (defined in Fig. 1) or log-power log-frequency slopes >-0.59 | PREP's 'findNoisyChannels', then for each 1s epoch, reject if >5% of data show extreme values (defined in Fig. 1) or log-power log-frequency slopes >-0.59 | PREP's 'findNoisyChannels', then for each 1s epoch, reject if >5% of data show extreme values (defined in Fig. 1) or log-power log-frequency slopes >-0.59 |
| **Initial Outlying Data Period Rejection** | After bad channels are rejected, mark remaining 1s periods that exceed the same thresholds as per the bad channel rejection step for exclusion from the MWF cleaning template and rejection prior to wICA. | After bad channels are rejected, reject remaining 1s periods that exceed the same thresholds as per the bad channel rejection step | After bad channels are rejected, reject remaining 1s periods that exceed the same thresholds as per the bad channel rejection step | After bad channels are rejected, mark remaining 1s periods that exceed the same thresholds as per the bad channel rejection step for exclusion from the MWF cleaning template and rejection after cleaning. |
| **Initial Artifact Reduction** | 3 sequential MWF runs, cleaning muscle activity first, then eye blinks, then horizontal eye movement and drift | None | None | 3 sequential MWF runs, cleaning muscle activity first, then eye blinks, then horizontal eye movement and drift |
| **Second Artifact Reduction** | ICA computed using the PICARD algorithm. Artifacts identified using Adjusted-ADJUST. Artifactual ICA components reduced using wICA. | ICA computed using PICARD. Artifactual ICA components subtracted (identified using Adjusted-ADJUST) | ICA computed using PICARD. Artifactual ICA components identified by Adjusted-ADJUST. Artifacts reduced using wICA. | None |

*Note.* Abbreviations: Hz = hertz; ICA = independent component analysis; PICARD = Preconditioned ICA for Real Data; PREP = Electroencephalography Pre-processing Pipeline; s = seconds; MWF = Multi-channel Wiener filters; wICA = wavelet enhanced independent component analysis.

### Comparison Pipelines

The comparison pipelines we tested were: i) the Maryland analysis of developmental EEG (MADE) pipeline (Debnath et al., 2020), ii) the Automated Pipeline for Infants Continuous EEG (APICE) (Flo et al., 2022), iii) the Harvard Automated Processing Pipeline for Electroencephalography (HAPPE) (Gabard-Durnam et al., 2018), and the Artifact Subspace Reconstruction approach followed by ICA subtraction of artifacts identified by Adjusted-ADJUST (referred to as ASR) (Chang et al., 2020).

### Segmentation of the Cleaned Data

After cleaning with each of the pipelines, EEG channels that were rejected during the cleaning steps were interpolated using spherical interpolation and data were segmented into 2-second non-overlapping epochs. Epochs rejection thresholds were applied as per the default RELAX settings (i.e., single/all channel improbable data thresholds [SD]: 5 and 3, respectively; single/all channel improbable data thresholds [SD]; 5 and 3, respectively; log-frequency log-power slope threshold for detecting muscle activity: -0.31), except for absolute voltage amplitude thresholds, which were set slightly more conservatively (+/- 100 uV) to avoid removing any epochs containing high-amplitude alpha activity.

### Cleaning Quality Evaluation Metrics

#### Multi-artifact Cleaning Performance Indicators: SER and ARR

Cleaning quality metrics used previously by our group (Bailey et al., 2023a; Bailey et al., 2023b) and others (Bertrand, 2015; Somers & Bertrand, 2016; Somers et al., 2018) were employed to examine pre-processing performance by each of the pipelines. Two complimentary metrics; the Signal-to-Error Ratio (SER) and Artifact-to-Residue Ratio (ARR) were used to estimate cleaning efficacy and preservation of the signal, respectively. SER measures the signal remaining unchanged after cleaning EEG data within periods of the continuous EEG data that were initially identified as free from all artifacts prior to cleaning. Artifact types included in the artifact templates included muscle activity, blinks, horizontal eye movements, and voltage drift. To understand how clean and contaminated periods were determined, refer to Bailey et al. (2023a). The SER measure is derived by dividing the expected value of the squared signal amplitude in clean periods for each electrode in the original (unprocessed) data by the squared signal of the removed artifacts during clean periods in the MWF templates (Somers et al., 2018). An average is then taken across all electrodes, with weighting applied according to the amplitude of the artifact signal in each electrode proportional to the amplitude across all electrodes. This results in electrodes containing greater artifact contributing more to the SER score. Larger SER values indicate better cleaning performance. ARR was calculated by obtaining the expected value operator of the square of the signal removed by the artifact reduction processes, divided by the expected value operator of the square of the raw data from the periods defined as containing artifacts prior to cleaning, minus the removed artifact signal from these artifact contaminated periods (Somers et al., 2018). As with the SER, larger ARR values are indicative of better cleaning performance. Further, given the complementary nature of these metrics, to perform well, pre-processing pipelines should be expected to achieve both high SER and ARR values (it is trivially easy but also unhelpful to clean artifacts very effectively if we are not concerned about also preserving the non-artifact signal).

#### Eye Blinks

Blink-Amplitude-Ratio (BAR) measures were also used to examine the ratio of blink amplitude to data periods containing no blinks (Robbins et al., 2020). BAR was assessed both across frontal channels (fBAR; average across channels: Fp1, Fp2, F9, F10, AF3, AF4; i.e., where blink amplitudes are typically maximal), as well as across all electrodes (allBAR). For these BAR measures, values close to 1 indicate optimal performance, with values below 1 indicating overcleaning and values larger than 1 indicating under cleaning (Robbins et al., 2020). Blink metrics were not assessed for the eyes-closed recordings, and 35 files were omitted from the blink metric analyses because there were not enough blink-specific segments of the EEG data left after removing epochs with multiple blinks.

#### Electromyographic Contamination

In order to provide an indication of how many epochs contained residual electromyographic (EMG) contamination after cleaning, we determined the number of EEG epochs with any electrode showing a log-power log-frequency slopes greater than -0.59, which previous research has indicated to reflect muscle activity (Fitzgibbon et al., 2016). Higher values represent poorer cleaning for this metric.

#### Proportion of Epochs Rejected

An important consideration for data cleaning is the number of data segments (epochs) that remain after pre- processing. It is typically beneficial to retain as much good quality data as possible. This can be particularly important in cases where, for example, recording durations are relatively short, as can be the case with EEG collected in neurodevelopmental cohorts who may only be able to remain still for brief periods. Lower values for the proportion of epochs rejected metric (indicating a smaller proportion of epochs rejected) reflects better performance.

#### Alpha Power and Alpha Peak Detection

As a final assessment, we investigated i) differences in alpha power values between the eyes-open and eyes-closed recordings, and ii) the likelihood of detecting an alpha peak in the data following cleaning. To achieve this, we first parameterised the data into periodic (oscillatory) and aperiodic components using the fitting oscillations and one over f (FOOOF) algorithm (Donoghue et al., 2020). We then calculated the peak alpha spectral power for each participant (i.e., the detected peak with the highest power within the 7-13 Hz range) (Donoghue et al., 2020). This metric provides a test of the potential for the cleaning algorithms to reveal experimental effects, using a well-established between condition comparison where a reduction in alpha power with eye-opening is observed, i.e., the ‘Berger Effect’ (Kirschfeld, 2005). We also further examined the percentage of total electrodes that had a detectable alpha peak following spectral parameterisation for each of the pipelines. As alpha is the dominant cortical rhythm at rest, and alpha oscillations are ubiquitous in resting-state neural recordings across much of the cortex (although most pronounced in posterior regions) (Edgar et al., 2023; Lew et al., 2021), we used this approach to assess the ability to detect alpha oscillations within the periodic EEG signal following processing with each of the pipelines.

### Statistical Analysis

Statistical analyses were performed in R (version 4.0.3) (R Core Team, 2020) and JASP (JASP Team, 2023). Robust repeated measures ANOVAs based on trimmed means were used to compare each of the pre-processing pipelines on the various cleaning quality evaluation metrics using the WRS2 package (Mair & Wilcox, 2020). This approach is robust to violations of normality and homoscedasticity, while maintaining equivalent power to traditional parametric tests (Mair & Wilcox, 2020). Significant omnibus ANOVAs were followed-up with pairwise comparisons using robust post-hoc t-tests, which apply Hochberg’s method to control for family-wise error (Hochberg, 1988; Mair & Wilcox, 2020). We did not perform additional experiment-wise multiple comparison controls (i.e., for each quality evaluation metric), to emphasize sensitivity for the detection of differences in cleaning outcomes, in alignment with our aim to assess the potential superiority of specific cleaning pipelines (Bailey et al., 2023a; Bailey et al., 2023b; Bender & Lange, 2001). We have also provided rank orders of the means for each pre-processing pipeline for each of the cleaning metrics to enable the reader to compare average cleaning performance for each metric across the pipelines (ranked from best performance to worst performance; Table 2). Finally, we have provided scatterplots of the SER x ARR values for each pre-processing pipeline to enable evaluation of these two complimentary metrics together (Figure 3). Heatmaps of the post-hoc tests for each metric are also provided in the Supplemental Materials (Figures S12-17). Bayesian analyses were also conducted to examine the strength of evidence supporting a power difference between eyes-open and eyes-closed recordings following cleaning by each pipeline.

**Figure 1:**
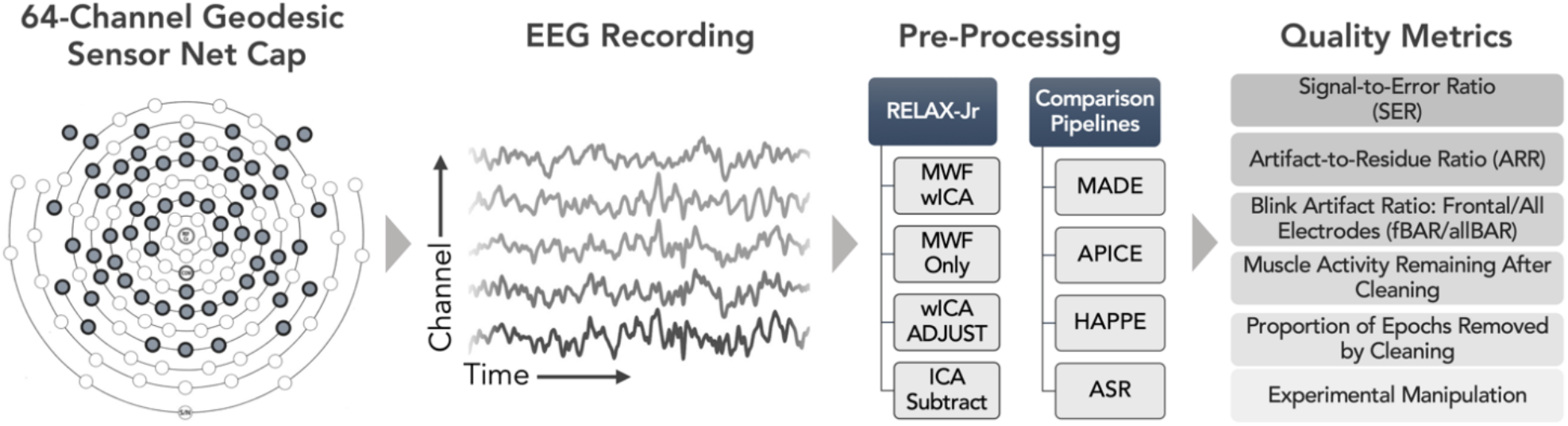
Overview of the data analysis pipeline. Eyes-open and eyes-closed resting-state EEG recordings were processed using four separate versions the RELAX-Jr pipeline (MWF wICA, MWF Only, wICA ADJUST, ICA Subtract), or using one of four comparison pipelines (MADE, APICE, HAPPE, ASR). Quality metrics were then used to assess data cleaning performance.

**Table 2.**
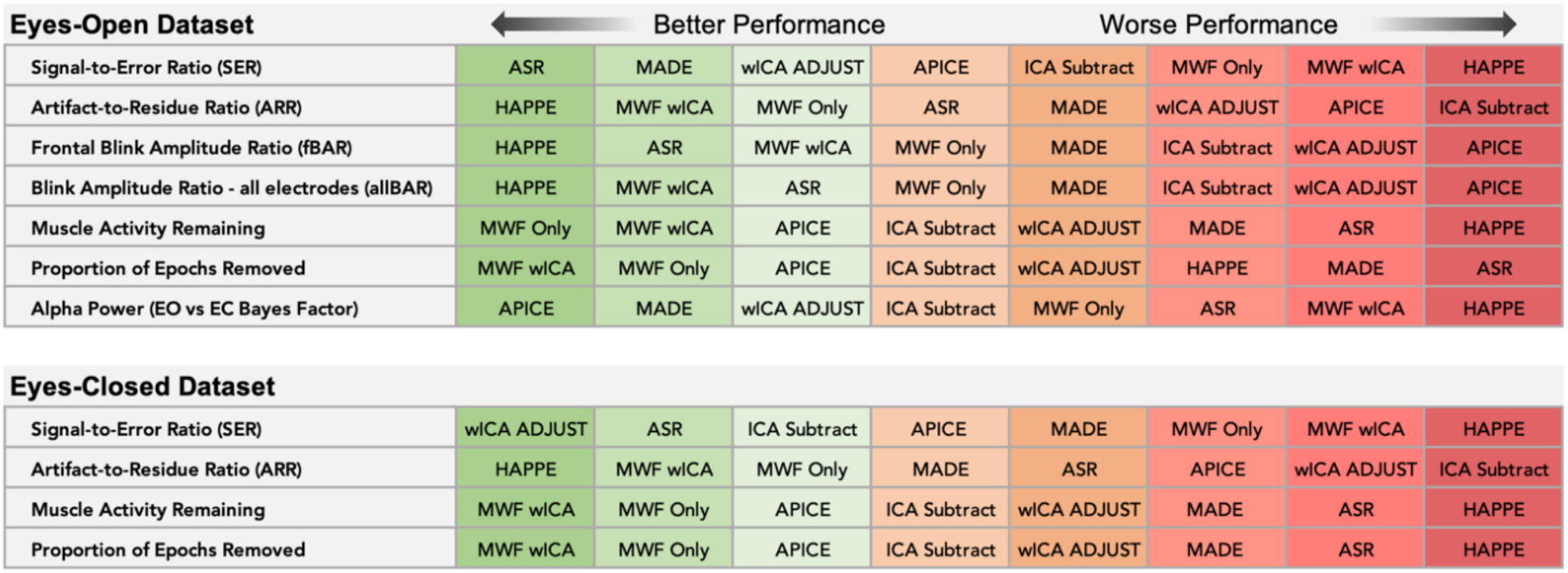
Rank order of each pipeline from best to worst performance (by mean). Note, either higher or lower values can reflect better performance (depending on the specific metric). In all instances, order is based on performance rather than value. Also note that blink-related metrics were only obtained for eyes-open datasets, while the alpha power metric was calculated as a percentage difference in power between the eyes-open and eyes-closed datasets.

| Eyes-Open Dataset |  | ← Better Performance |  |  | Worse Performance → |  |  |  |
| --- | --- | --- | --- | --- | --- | --- | --- | --- |
| Signal-to-Error Ratio (SER) | ASR | MADE | wICA ADJUST | APICE | ICA Subtract | MWF Only | MWF wICA | HAPPE |
| Artifact-to-Residue Ratio (ARR) | HAPPE | MWF wICA | MWF Only | ASR | MADE | wICA ADJUST | APICE | ICA Subtract |
| Frontal Blink Amplitude Ratio (fBAR) | HAPPE | ASR | MWF wICA | MWF Only | MADE | ICA Subtract | wICA ADJUST | APICE |
| Blink Amplitude Ratio - all electrodes (allBAR) | HAPPE | MWF wICA | ASR | MWF Only | MADE | ICA Subtract | wICA ADJUST | APICE |
| Muscle Activity Remaining | MWF Only | MWF wICA | APICE | ICA Subtract | wICA ADJUST | MADE | ASR | HAPPE |
| Proportion of Epochs Removed | MWF wICA | MWF Only | APICE | ICA Subtract | wICA ADJUST | HAPPE | MADE | ASR |
| Alpha Power (EO vs EC Bayes Factor) | APICE | MADE | wICA ADJUST | ICA Subtract | MWF Only | ASR | MWF wICA | HAPPE |
| Eyes-Closed Dataset |  |  |  |  |  |  |  |  |
| Signal-to-Error Ratio (SER) | wICA ADJUST | ASR | ICA Subtract | APICE | MADE | MWF Only | MWF wICA | HAPPE |
| Artifact-to-Residue Ratio (ARR) | HAPPE | MWF wICA | MWF Only | MADE | ASR | APICE | wICA ADJUST | ICA Subtract |
| Muscle Activity Remaining | MWF wICA | MWF Only | APICE | ICA Subtract | wICA ADJUST | MADE | ASR | HAPPE |
| Proportion of Epochs Removed | MWF wICA | MWF Only | APICE | ICA Subtract | wICA ADJUST | MADE | ASR | HAPPE |

## RESULTS

The omnibus ANOVA was significant for all metrics tested for both the eyes-open and eyes-closed datasets (specific details in the sections below). A rank order of the means for each pre-processing pipeline is provided in Table 2. Due to the large number of comparisons across the various pipelines, we provide here a summary only of the most relevant results, with more detailed results from the follow-up post-hoc tests provided in the Supplemental Materials (Figures S12-17). An example of a raw (unprocessed) and cleaned EEG trace (single subject) is provided in Figure 6, while further examples for all pipelines are provided in the Supplemental Materials (Figures S3-11).

### Signal-to-Error Ratio and Artifact-to-Error Ratio

The omnibus ANOVA was significant for both the eyes-open, *F*(3.57, 278.78) = 83.58, *p* < 0.0001, and eyes-closed, *F*(3.71, 300.61) = 76.84, *p* < 0.0001, conditions, indicating a significant difference in SER values between the pipelines (see Figure 2 for plots of SER and ARR values). For the ARR, the omnibus ANOVA was significant for both the eyes-open, *F*(3.82, 298.35) = 417.06, *p* < 0.0001, and eyes-closed, *F*(3.61, 292.81) = 524.00, *p* < 0.0001, conditions. For both the eyes-open and eyes-closed data, when assessing the SER and ARR metrics together (Figure 3), the MWF wICA and MWF Only pipelines demonstrated a good middle-ground between both values, with moderate scores on both metrics. In contrast, the ASR, wICA ADJUST, MADE, APICE, and ICA Subtract pipelines all showed relatively high SER values, but had lower ARR values, indicating that although these pipelines produced little distortion or reduction of the clean EEG signal, they were generally less effective at suppressing artifact. The most extreme examples of this were the ASR pipeline for the eyes-open data, and the wICA ADJUST, ICA Subtract, and ASR pipelines for the eyes-closed data, which demonstrated very high SER and low ARR values. In contrast, the HAPPE pipeline demonstrated very high ARR and very low SER values in both the eyes open and closed datasets, indicating that while it was effective at mitigating artifacts, it concurrently removed a substantial amount of the signal from the clean segments of the EEG recordings.

**Figure 2:**
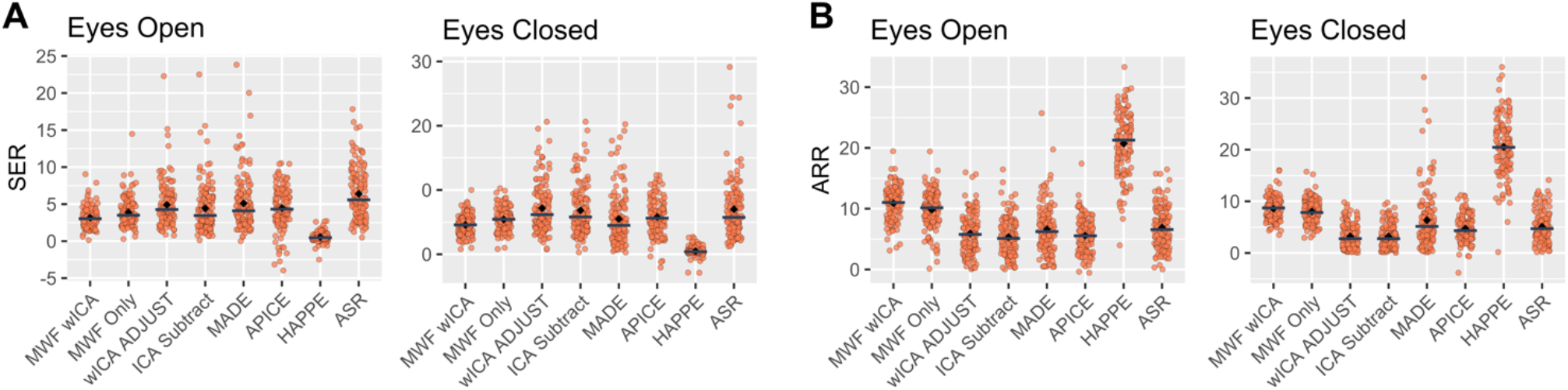
Plots of A) Signal-to-Error (SER) and B) Artifact-to-Residue (ARR) values for the eyes-open and eyes-closed recordings. Grey horizontal lines denote the median, while black diamonds denote the mean. Higher values for both SER and ARR concurrently reflect better cleaning performance (higher SER = better signal preservation; higher ARR = better artifact reduction).

**Figure 3:**
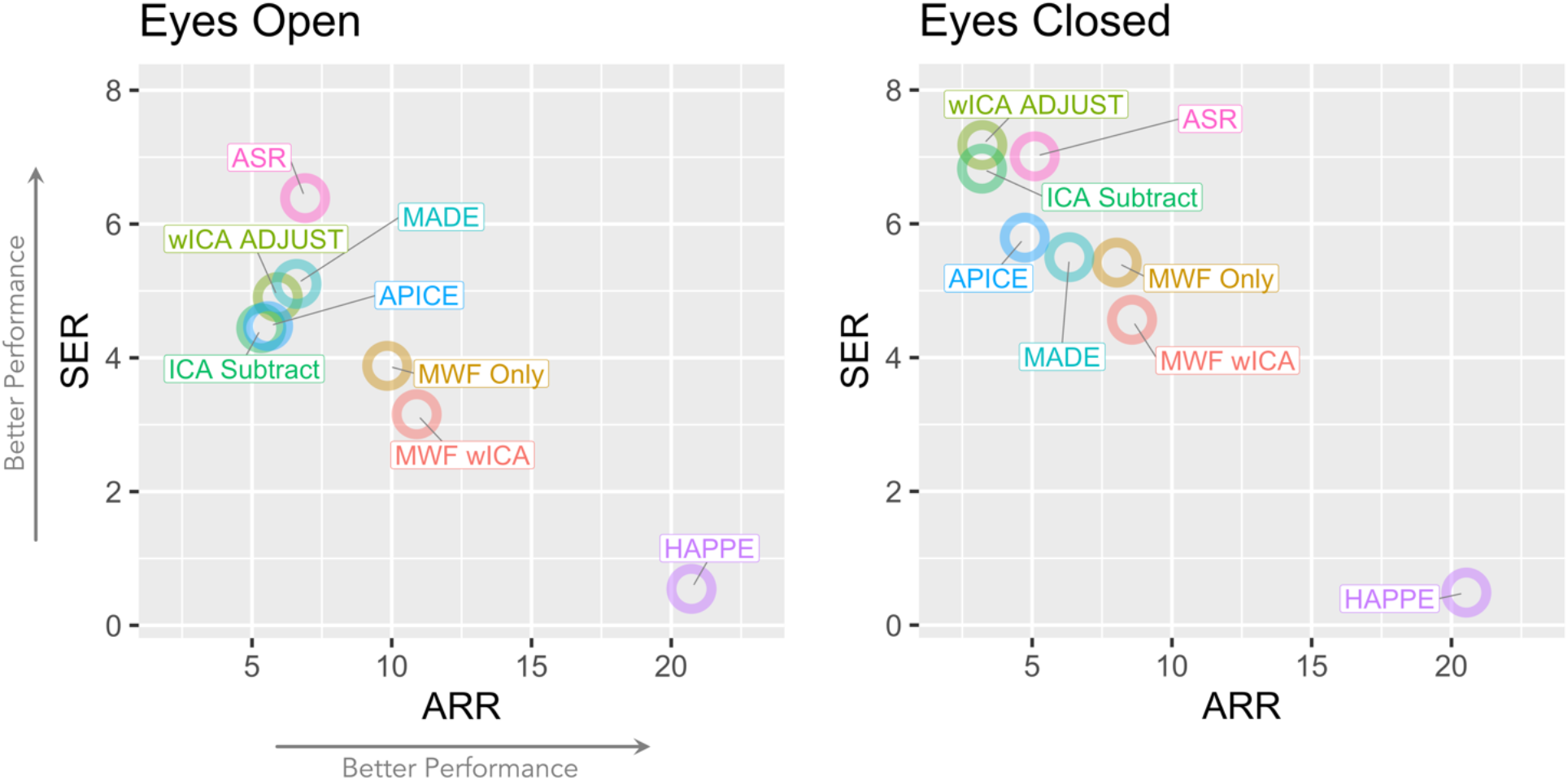
Scatterplots depicting the mean Signal-to-Error (SER) and Artifact-to-Residue (ARR) values for the eyes-open and eyes-closed recordings for each of the pre-processing pipelines. Higher SER and ARR values reflect better cleaning of the EEG data.

It is also interesting to note that for the eyes-open data, the ASR pipeline resulted in higher values for SER but similar values for ARR compared to ICA Subtract, despite including ICA Subtract as one of the cleaning steps in the ASR pipeline. In contrast, MWF wICA showed higher ARR but lower SER than both pipelines that represented components of the combined MWF wICA pipeline (MWF Only and wICA ADJUST). One possible mechanism by which ASR (which combines ASR and ICA Subtract) could produce increased SER compared to ICA Subtract only is that the initial artifact reduction performed by ASR allowed the ICA Subtract step to better separate artifact components from neural components, better preserving the clean signal. However, MWF wICA applies an analogous approach, first reducing artifacts with MWF prior to wICA, so if this mechanism were accurate, then we might expect MWF wICA and wICA ADJUST to show the same pattern. As such, another potential mechanism worth considering is that ASR may enhance the amplitude of activity in the clean periods of the data, leading to higher SER values through an artificial increase in amplitude in clean periods rather than the preservation of signal in those clean periods.

### Blink Amplitude Ratio

For the fBAR values taken from anterior electrodes in close proximity to the eyes, the omnibus ANOVA was significant *F*(3.03, 154.63) = 59.80, *p* < 0.0001. For the allBAR values (all electrodes), the ANOVA was also significant, *F*(3.32, 169.54) = 111.56, *p* < 0.0001. The HAPPE pipeline performed the best across the fBAR and allBAR metrics, followed by ASR for fBAR and MWF wICA for allBAR, with APICE performing the worst for both the fBAR and allBAR metrics (values closest to 1 = best performance; see Table 2 for rank orders for each pipeline, and Table 3 for Mean and SD values). BAR values are depicted in Figure 4.

**Figure 4:**
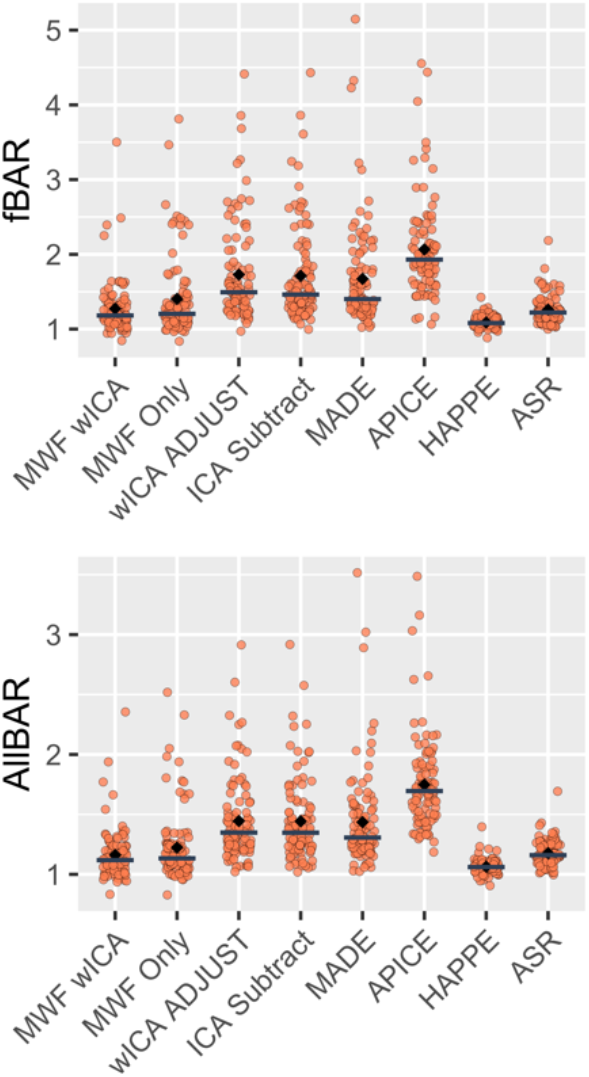
Blink amplitude ratios across frontal electrodes (average across Fp1, Fp2, F9, F10, AF3, AF4; fBAR) and across all electrodes (allBAR). Grey horizontal lines denote the median while black diamonds denote the mean.

**Table 3:** Means and standard deviations (in parentheses) of the quality evaluation metric values for the eyes-open data. Higher values for SER and ARR indicate better cleaning performance. For fBAR and allBAR, values ∼1 indicate optimal cleaning performance, while for EMG remaining (i.e., proportion of epochs showing EMG after cleaning) and epochs removed (proportion of epochs removed by cleaning), lower values indicate better performance.

| Pipeline | SER | ARR | fBAR | allBAR | EMG Remaining | Epochs Removed |
| --- | --- | --- | --- | --- | --- | --- |
| MWF wICA | 3.16 (1.37) | 10.89 (2.72) | 1.27 (0.35) | 1.16 (0.21) | 0.003 (0.010) | 0.07 (0.03) |
| MWF Only | 3.88 (1.90) | 9.84 (3.06) | 1.40 (0.52) | 1.22 (0.29) | 0.003 (0.009) | 0.09 (0.05) |
| wICA ADJUST | 4.91 (2.95) | 5.92 (3.32) | 1.73 (0.65) | 1.45 (0.35) | 0.02 (0.045) | 0.15 (0.12) |
| ICA Subtract | 4.44 (3.24) | 5.33 (3.17) | 1.71 (0.64) | 1.44 (0.34) | 0.019 (0.044) | 0.15 (0.13) |
| MADE | 5.10 (3.73) | 6.59 (4.01) | 1.67 (0.67) | 1.44 (0.39) | 0.037 (0.059) | 0.19 (0.15) |
| APICE | 4.47 (2.79) | 5.58 (2.88) | 2.07 (0.63) | 1.75 (0.39) | 0.015 (0.037) | 0.12 (0.10) |
| HAPPE | 0.55 (0.62) | 20.72 (5.17) | 1.09 (0.08) | 1.06 (0.06) | 0.988 (0.044) | 0.17 (0.31) |
| ASR | 6.39 (3.53) | 6.90 (3.46) | 1.26 (0.19) | 1.18 (0.11) | 0.097 (0.097) | 0.25 (0.21) |

### Proportion of Epochs Showing Muscle Activity After Cleaning

The omnibus ANOVA was significant for both the eyes-open, *F*(1.2, 93.67) = 17231.98, *p* < 0.0001, and eyes-closed, *F*(2.08, 168.17) = 48712.90, *p* < 0.0001, conditions. MWF wICA and MWF Only were the best performers in terms of the proportion of epochs showing EMG after cleaning, with MWF Only showing the best overall performance for the eyes-open dataset, and MWF wICA showing the best performance for the eyes-closed data. For both pipelines, the amount of EMG remaining was extremely close to zero, and means were identical to three decimal places (see Tables 3 and 4 for Means and SDs). The order of performance was the same for the rest of the pipelines for both the eyes-open and eyes-closed datasets, with the next best being APICE, followed by ICA Subtract, wICA ADJUST, MADE, ASR, and finally HAPPE (see Figure 5A). The HAPPE pipeline showed notably poor performance on this metric, with the majority of epochs showing log-power log-frequency slopes that exceeded thresholds for indicating residual EMG following cleaning.

**Figure 5:**
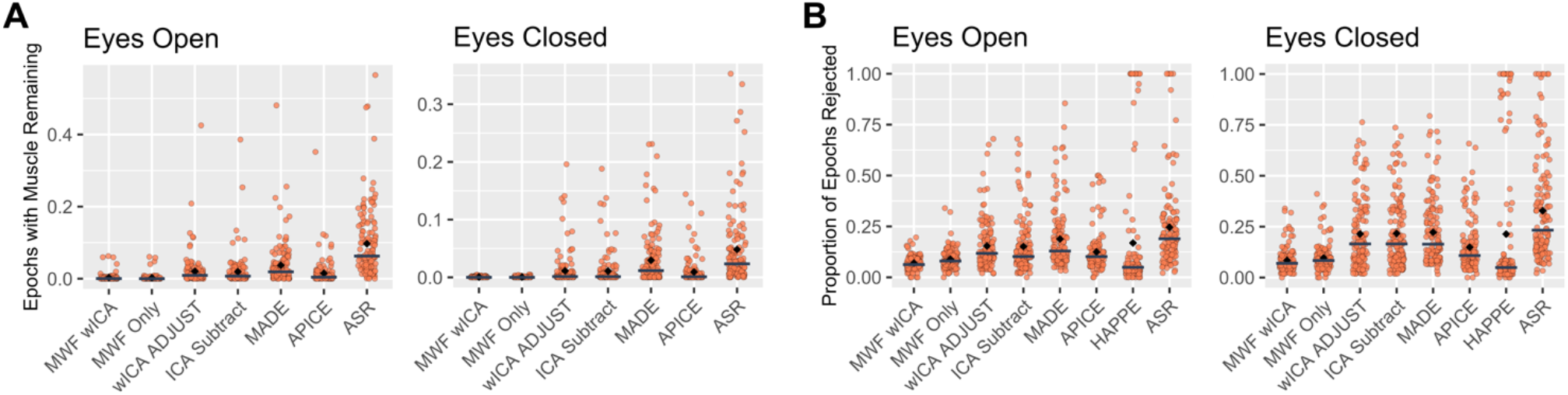
A) Proportion of epochs showing log-power log-frequency values above the -0.59 threshold for EMG activity remaining for each of the pre-processing pipelines. Lower values reflect more effective cleaning of EMG activity. Grey horizontal lines denote the median while black diamonds denote the mean. Note the data presented here were first winsorised (z = +/-2.5) as outlying values from the MADE and ASR pipelines made it difficult to visualise differences between the pipelines. The HAPPE pipeline was also removed as it had a median value >0.95. Plots of the full dataset can be found in the Supplemental Materials (Figure S1). B) Proportion of epochs rejected after cleaning for each pipeline.

**Figure 6:**
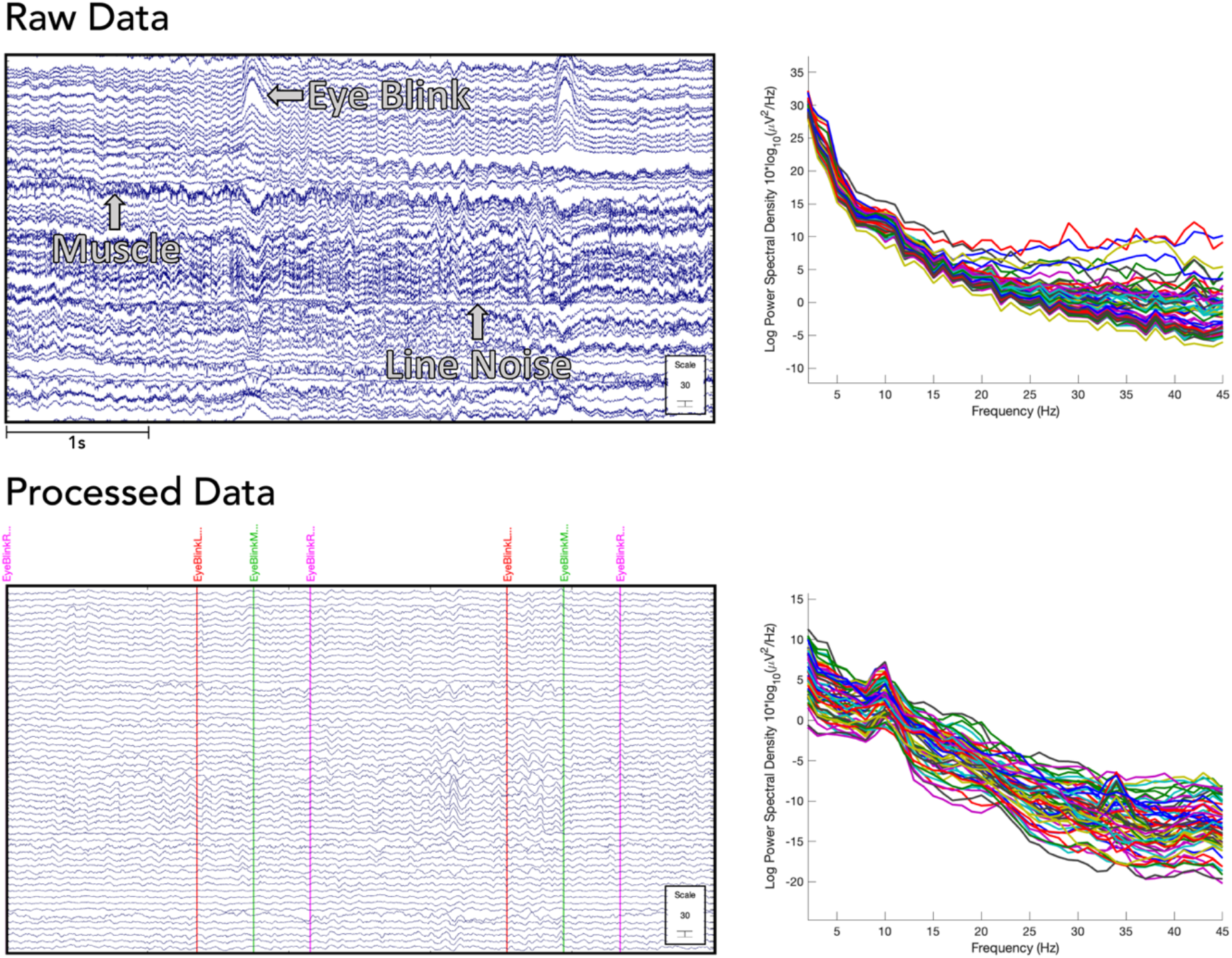
Left: EEG signal from a single example subject (eyes-open recording, 5 second segment) showing the raw data (with DC offset removed [top]) and following pre-processing with RELAX-Jr (MWF wICA pipeline [bottom]). Examples of several common artifacts are noted on the EEG trace. All scales are microvolts. Right: Power spectra from the same subject before (top) and after (bottom) cleaning with the same RELAX-Jr pipeline (power spectra are from the entire 2 min recording).

**Table 4:** Means and standard deviations (in parentheses) of the quality evaluation metric values for the eyes-closed data. For EMG remaining (i.e., proportion of epochs showing EMG after cleaning) and epochs removed (proportion of epochs removed by cleaning), lower values indicate better performance.

| Pipeline | SER | ARR | EMG Remaining | Epochs Removed |
| --- | --- | --- | --- | --- |
| MWF wICA | 4.57 (1.56) | 8.57 (2.27) | 0.000 (0.001) | 0.08 (0.06) |
| MWF Only | 5.42 (1.69) | 8.03 (2.24) | 0.000 (0.001) | 0.10 (0.07) |
| wICA ADJUST | 7.18 (3.88) | 3.22 (2.37) | 0.011 (0.029) | 0.21 (0.17) |
| ICA Subtract | 6.82 (3.89) | 3.20 (2.36) | 0.011 (0.028) | 0.22 (0.17) |
| MADE | 5.50 (4.12) | 6.33 (5.58) | 0.030 (0.046) | 0.22 (0.16) |
| APICE | 5.79 (2.73) | 4.73 (2.51) | 0.009 (0.023) | 0.15 (0.13) |
| HAPPE | 0.49 (0.70) | 20.53 (5.71) | 0.968 (0.075) | 0.21 (0.34) |
| ASR | 7.01 (4.73) | 5.09 (3.04) | 0.048 (0.066) | 0.33 (0.25) |

### Proportion of Epochs Removed by Cleaning

The omnibus ANOVA was significant for both the eyes-open, *F*(4.21, 328.75) = 53.70, *p* < 0.0001, and eyes- closed, *F*(3.51, 284.08) = 38.39, *p* < 0.0001, conditions. For both the eyes-open and eyes-closed data, MWF wICA performed the best, followed by MWF Only, both of which resulted in the rejection of only 10% of the data or less on average, with MWF wICA resulting in the preservation of more than 80% of epochs for all EEG files in the eyes open data. In contrast, the worst performers (in terms of mean values) in the eyes open data were MADE, ASR and HAPPE, with ASR and HAPPE pipelines rejecting 100% of epochs for some files, and ASR rejecting more than 25% of epochs on average. In the eyes closed data, wICA ADJUST, ICA Subtract, MADE, and HAPPE all rejected more than 20% of epochs on average, ASR rejected 33% of epochs, and both ASR and HAPPE rejected 100% of epochs for some EEG files (see Figure 5B).

### Variance Explained by Experimental Manipulation (Berger effect)

Figure 7 shows the frequency spectra of the oscillatory neural activity (i.e., after removal of the aperiodic signal) for both the eyes-open and eyes-closed datasets. This figure highlights the alpha band demonstrating the distribution of alpha power across the scalp, while Figure 8 depicts the individual differences between the datasets for each pipeline using a power value derived from all electrodes (root mean square [RMS]). All pipelines showed the expected pattern of enhanced alpha power during eye closure, relative to the eyes-open condition. To statistically assess differences between the eyes-open and eyes-closed conditions in terms of the strength of alpha activity differences between eyes-open and eyes-closed conditions, a Bayesian approach was used to determine the strength of the evidence supporting a power difference between the two conditions. Specifically, RMS alpha power values were compared between the eyes-open and eyes-closed conditions using Bayesian paired-samples t-tests. Bayes factors provided extremely strong support for the alternative hypothesis for all pipelines, indicating an expected pattern of robust alpha power differences between the eyes-closed and eyes-open conditions (see Supplemental Table S1 for complete table of results). APICE showed the highest Bayes factor value, followed by MADE, wICA ADJUST, ICA Subtract, MWF Only, ASR, and MWF wICA, with HAPPE showing the lowest Bayes factor value. Next, we computed difference scores ([eyes-closed] – [eyes-open]) for RMS power values for each individual participant after cleaning with each pipeline. We used these difference scores to statistically compare each of the pipelines using a one-way ANOVA. The omnibus ANOVA was significant, *F*(7,1059) = 8.952, *p* < 0.001; however, post-hoc tests (Bonferroni corrected) only revealed significant differences between the HAPPE pipeline and all other pipelines (all p < 0.001, with HAPPE showing a smaller difference than all other pipelines), with none of the other pipelines differing in terms of difference scores between eyes-open and eyes-closed power.

**Figure 7:**
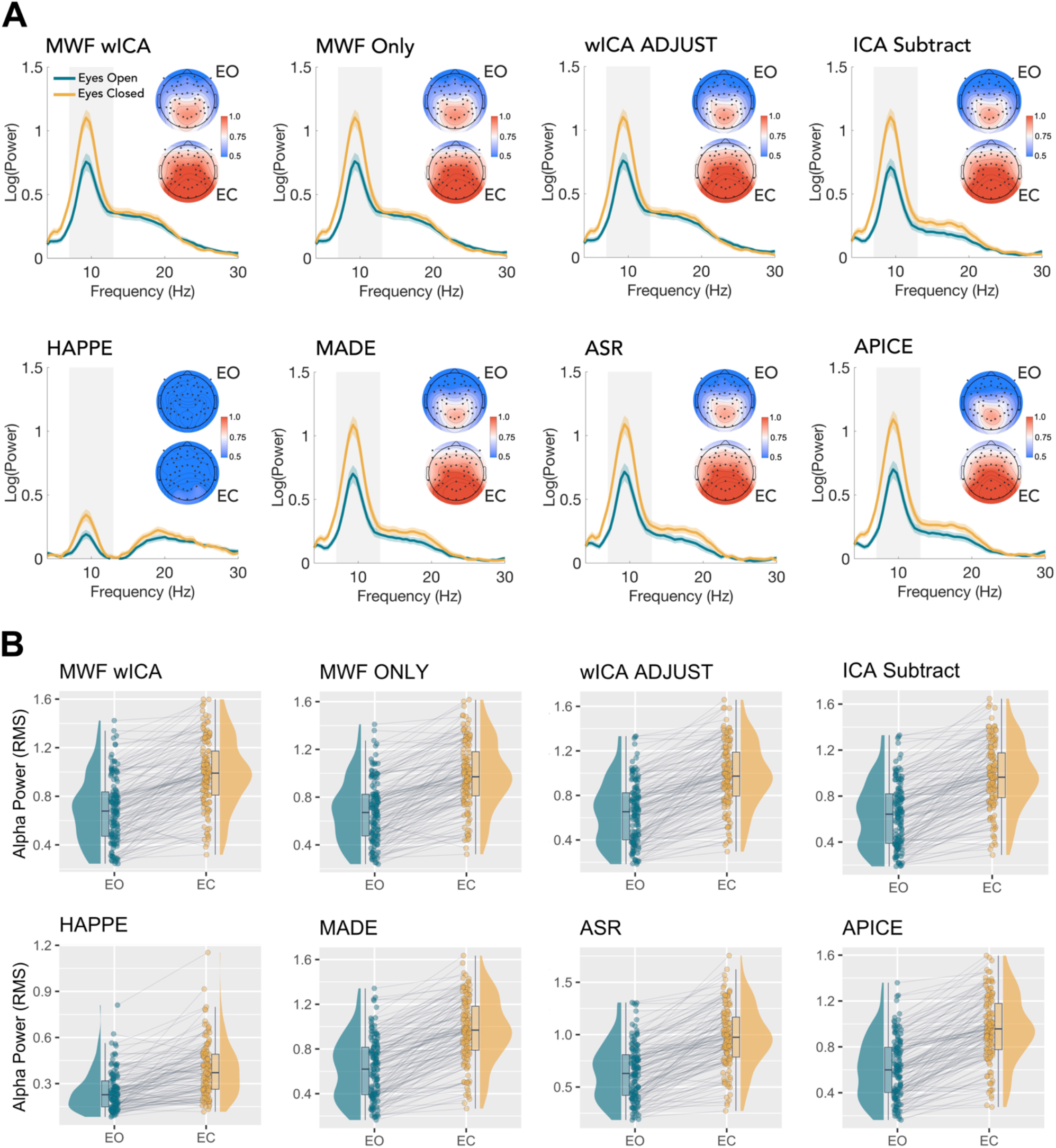
A) Power spectral density plots (POz electrode) of the eyes-open and eyes-closed data after removal of the aperiodic signal. Shaded line is 95% confidence interval. Translucent grey bar denotes the alpha frequency range (7-13 Hz). Accompanying topographic plots depict the distribution of alpha power across the scalp. B) Raincloud plots showing differences in alpha power (root mean square [RMS] values) between the eyes-open (EO) and eyes-closed (EC) conditions for each of the pipelines.

**Figure 8:**
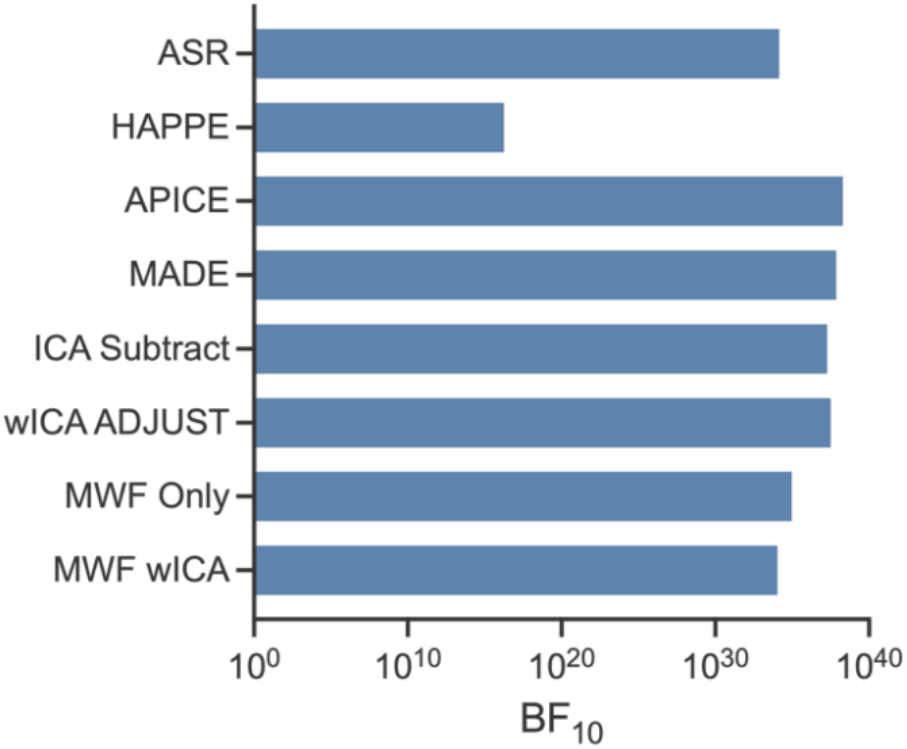
Plots of BF10 values for the Bayesian t-tests comparing the eyes-open and eyes-closed RMS alpha power values. Note, x-axis is log-scale to improve visualisation across results for the different pipelines which had large disparity.

### Alpha Peak Detection

Separate one-way ANOVAs were run for the eyes-open and eyes-closed recording conditions comparing each pipeline in terms of the percentage of total electrodes that had a detectable alpha peak following spectral parameterisation with the FOOOF algorithm to account for the aperiodic signal (Figure 9). The omnibus ANOVAs were significant for both conditions (eyes open: *F*(7,1037) = 41.155, *p* < 0.001; eyes-closed: *F*(7,1062) = 22.898, *p* < 0.001). For both conditions, Bonferroni corrected post-hoc tests indicated that the HAPPE pipeline had a significantly lower percentage of electrodes with a detected alpha peak compared to all other pipelines (all p < 0.001). No significant differences were observed between any of the other pipelines. Additional plots comparing pipelines across theta (4-7 Hz) and beta (13-30 Hz) peaks are also available in the Supplementary Material (Figure S2).

**Figure 9:**
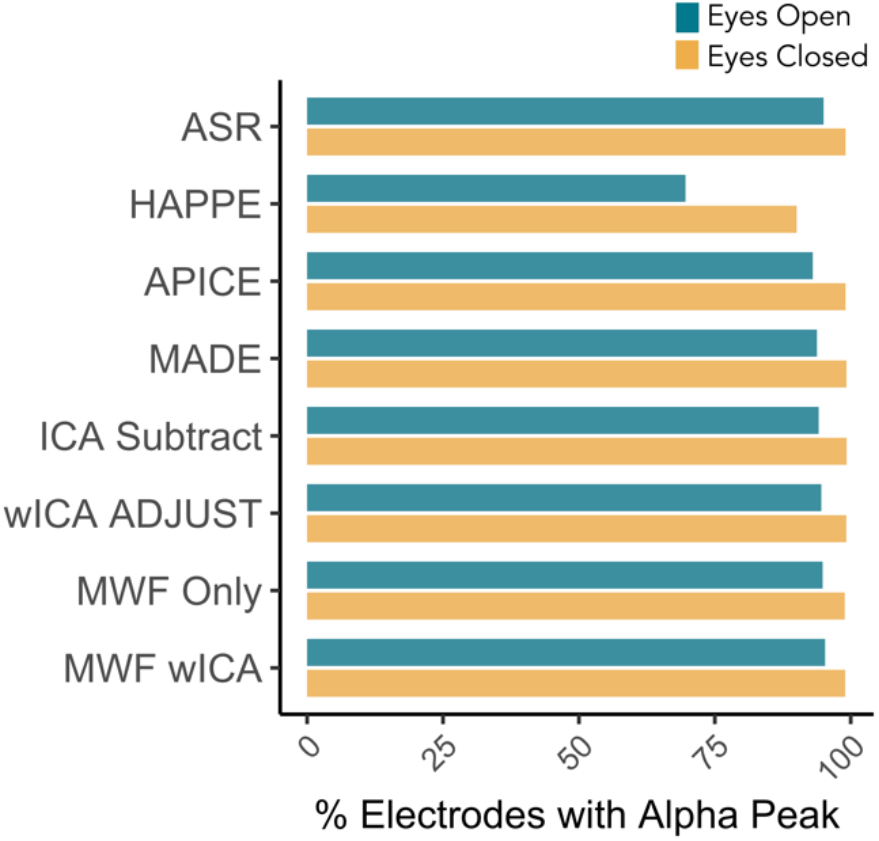
Percentage of total electrodes having a detected peak within the alpha band after removal of the 1/*f*-like aperiodic signal for the eyes-open and eyes-closed data after cleaning with each pipeline (average over all participants).

## Discussion

This study sought to provide an overview and assessment of the newly developed RELAX-Jr pre-processing pipeline for use with EEG data recorded in children. This software provides an important extension to the original RELAX software, which was designed for, and extensively evaluated in, adult populations (Bailey et al., 2023a; Bailey et al., 2023b). We have optimised this pipeline through the inclusion of the PICARD ICA algorithm, which has been demonstrated to perform equivalently to extended-infomax ICA but is computationally much faster (Ablin et al., 2018a), as well as the adjusted-ADJUST independent component classification algorithm specifically designed for use with data collected from children. This allows RELAX-Jr to provide a comprehensive integrated MATLAB-based approach for cleaning data collected from neurodevelopmental cohorts, which, in addition to the neural signal of interest, often also contain high levels of artifact. We tested four versions of the RELAX-Jr software on resting-state EEG data collected in children aged between 4-to-12 years of age. We compared our software against four popular open-source automated EEG cleaning pipelines. Results from our comprehensive assessment indicate that RELAX-Jr generally performed well overall across the majority of included metrics, in particular the MWF wICA and MWF Only settings providing superior performance in terms of preserving epochs for analysis and reducing the impact of muscle artifacts on the data, while also providing amongst the highest performance for blink artifact reduction (only being outperformed by HAPPE, which may over-clean the data) and statistically similar detection of the Berger effect compared to other pipelines.

The MWF wICA and MWF Only versions of the pipeline demonstrated strong overall performance when considering all metrics. The SER and ARR metrics indicated that these pipelines performed well in terms of removing artifacts from the data, while preserving the neural signal. In contrast, while the ICA Subtract and wICA ADJUST demonstrated higher SER values indicating that they were able to preserve more of the signal, they were less effective at removing artifact, as indicated by lower ARR values. This was most apparent for the eyes-closed recordings, where two pipelines obtained amongst the highest scores for SER, but the lowest scores for ARR. In contrast, the very high ARR score combined with very low SER score obtained by the HAPPE pipeline suggests that it was likely to be overcleaning the data. We also note that ASR performed strongly (i.e., ASR showed the best performance for SER), indicative of its strong cleaning performance. The MWF Only and MWF wICA pipelines also performed exceptionally well in terms of successful removal of EMG activity from the EEG recordings and minimising the number of artifact-contaminated epochs meeting the threshold for removal after cleaning. MWF wICA and MWF Only also showed statistically equivalent performance to the second-best performing pipeline (ASR) in terms of blink artifact reduction. In contrast, APICE performed considerably worse than other pipelines, with an average fBAR value of >2 indicating that blink periods in the data were twice the amplitude of non-blink periods following cleaning, thus suggesting inadequate removal of blinks from the data. We note that while the HAPPE pipeline performed the best for blink artifact reduction, the very low SER values observed, as well as very low amplitude EEG activity following cleaning by HAPPE, suggest that HAPPE’s excellent blink reduction performance likely comes at the expense of excessive over-cleaning of the EEG recording, where considerable neural signal is also removed from the data. This closely mirrored the very low amplitude recordings obtained after pre-processing with HAPPE when comparisons were made to the RELAX pipeline in adults (Bailey et al., 2023a).

It may be interesting to consider the contrast to our conclusions for the optimum pipeline in adult data, where MWF wICA or wICA ICLabel were recommended as default pipelines (with wICA ICLabel recommended when a sufficient amount of data are available, as it produced higher effect sizes for task related frequency band power experimental outcomes despite inferior cleaning to MWF wICA). In the current study, although wICA ADJUST was adapted to optimize its application to childhood EEG recordings, MWF Only performed better, despite not being specifically adapted to childhood EEG recordings. It may be that wICA ADJUST was not able to sufficiently clean the childhood EEG data due to the increased frequency and severity of artifacts, allowing MWF Only and MWF wICA to show higher performance. Future research may be able to optimize MWF Only for application to childhood EEG recordings by systematically testing the thresholds used to identify blink, muscle, and horizontal eye movement artifacts. However, we note that the muscle artifact thresholds used within RELAX were identified through research that involved paralysing participant’s scalp muscles prior to EEG to eliminate the potential for scalp EMG (Fitzgibbon et al., 2016), an approach that is unlikely to be feasible in children.

In addition to our results for artifact cleaning, it was also encouraging to see that all cleaning pipelines enabled clear differentiation between the eyes-open and eyes-closed recordings, indicating sensitivity to this experimental manipulation. Specifically, all pipelines revealed the expected finding of higher alpha power during the eyes-closed relative to eyes-open recordings, with Bayes factors showing extremely strong evidence in favour of the alternative hypothesis. Further, when comparing each of the pipelines directly (using difference scores between the [eyes-closed] – [eyes-open] RMS power values), there were no significant differences between the pipelines in their ability to detect the Berger effect (with the exception of HAPPE, which showed lower difference values compared to all other pipelines). Similarly, with the exception of HAPPE, the pipelines did not differ in terms of the percentage of electrodes with a detected alpha peak following removal of the aperiodic signal. This suggests that experimental outcomes are likely to be similar regardless of the cleaning pipeline used, so decisions about which pipeline to implement may be made based on which pipelines provide the best artifact reduction (as long as this is concurrent with sufficient preservation of the neural signal). This observation corroborates our previous results comparing pre-processing pipelines in adults (Bailey et al., 2023a). However, we note that the Berger effect (i.e., alpha blocking) is a robust phenomenon that typically produces large differences in EEG amplitude between conditions (Goncharova & Barlow, 1990; Kirschfeld, 2005; Niedermeyer, 1997). As such, it is possible that more subtle differences, such as disease-specific alterations in neural activity, or changes following therapeutic interventions, might be more heavily influenced by specific pre-processing pipelines. Future work comparing pre-processing pipelines across participants with various neurodevelopmental and/or neuropsychiatric diagnoses might be useful to explore this possibility.

## Limitations and Future Directions

We only tested RELAX-Jr on 64-channel Geodesic Sensor-Net EEG caps. As such, its effectiveness for cleaning data from higher density (e.g., 128, or 256 electrode) montages is uncertain. However, we do not foresee any obvious constraints that might prohibit applying RELAX-Jr to higher density recordings other than a likely increase in analysis times arising from greater computational burden produced by the inclusion of larger data files. The RELAX-Jr software was also assessed using a developmental dataset containing a relatively wide participant age range (4-12 years), and we did not assess its performance using data from infants or very young children. As EEG recordings in these populations can deviate substantially from older children and adults, including greater prevalence of low frequency activity (Hrachovy & Mizrahi, 2016; Marshall et al., 2002), it is possible that artifact cleaning performance in samples of infants might differ from that reported using the present cohort. Future work examining performance in younger cohorts and incorporating datasets with high-density recordings, would therefore be beneficial.

Additionally, as RELAX-Jr was assessed using a typically developing sample of children, its cleaning performance in clinical samples (e.g., autism, attention deficit hyperactivity disorder, epilepsy) remains to be established. We also note that lower density recordings, particularly those with <32 electrodes, could achieve significantly poorer performance resulting from insufficient ICA decomposition, which may not adequately separate neural from artifactual components (Janani et al., 2018; Klug & Gramann, 2021). For such recordings, the use of non-ICA based methods for data pre-processing should be considered. The efficacy of MWF for artifact reduction with <32 electrodes is currently untested, but we suspect it may still perform adequately for reducing artifacts in recordings with <32 electrodes. However, since the MWF approach acts as a spatial filter, its performance will have a lower boundary in terms of numbers of available electrodes, and we encourage future research to test where this lower boundary lies. We also note that recent automated pipelines have been developed specifically for data with low numbers of channels (e.g., HAPPILEE; Lopez et al., 2022) and may be beneficial for investigators analysing very low-density EEG recordings. We further observe that the Adjusted-ADJUST algorithm (Leach et al., 2020) integrated into the RELAX-Jr pipeline for objective selection of independent components representing artifacts requires the presence of left and right anterior electrodes for classification of blinks and horizontal eye movements. It is therefore possible that, in rare cases, a file with very high levels of artifact across frontal regions could lead to problems if large numbers of electrodes are removed. In such cases, more liberal extreme rejection thresholds could be considered, and might still provide adequate performance (Delorme, 2023). Finally, we note that our tests of experimental effects only include the examination of differences in the alpha frequency band, and that the current results may not apply to event-related potential analyses. In adult populations, our previous research with the RELAX software indicated that the wICA ICLabel setting was optimal for clean data where many epochs were available, and that MWF wICA might be preferred where data are noisier or fewer epochs are available. However, it is worth noting that more aggressive (e.g., 1 Hz) high pass filter settings are not appropriate for event-related potential analyses (as commonly analysed slow latency event-related potentials contain frequencies below 1Hz), but also that all artifact reduction pipelines performed more poorly in adult data when 0.25 Hz high pass filters (which are appropriate for event-related potential analyses) were implemented. As such, further research is required to determine the best approach for analysing event-related potentials in childhood EEG data.

## Conclusion

Here, we have provided an assessment of the cleaning performance of the RELAX-Jr software pipeline developed for pre-processing of EEG data collected from neurodevelopmental populations. The aim of this software is to provide users with a versatile and fully automated toolbox for removing artifacts that frequently contaminate the EEG record, enabling reliable and reproducible cleaning of EEG datasets while preserving the neural signal and minimising user bias (and workload). Based on the results of our analyses, we recommend the MWF wICA implementation of the software for applications to child EEG data, given its strong performance across the range of metrics assessed, including the ability to maximise the number of epochs included for analysis, which can be an important consideration for developmental EEG recordings, which are often limited in length.

## References

Ablin, P., Cardoso, J. F., & Gramfort, A. (2018a, 15-20 April 2018). Faster ICA Under Orthogonal Constraint. Paper presented at the 2018 IEEE International Conference on Acoustics, Speech and Signal Processing (ICASSP).

Ablin, P., Cardoso, J. F., & Gramfort, A. (2018b). Faster Independent Component Analysis by Preconditioning With Hessian Approximations. IEEE Transactions on Signal Processing, 66(15), 4040–4049. doi:10.1109/TSP.2018.2844203

Bailey, N. W., Biabani, M., Hill, A. T., Miljevic, A., Rogasch, N. C., McQueen, B., . . . Fitzgerald, P. B. (2023a). Introducing RELAX: An automated pre-processing pipeline for cleaning EEG data - Part 1: Algorithm and application to oscillations. Clin Neurophysiol. doi:10.1016/j.clinph.2023.01.017

Bailey, N. W., Hill, A. T., Biabani, M., Murphy, O. W., Rogasch, N. C., McQueen, B., . . . Fitzgerald, P. B. (2023b). RELAX part 2: A fully automated EEG data cleaning algorithm that is applicable to Event-Related-Potentials. Clin Neurophysiol. doi:10.1016/j.clinph.2023.01.018

Bell, A. J., & Sejnowski, T. J. (1995). An information-maximization approach to blind separation and blind deconvolution. Neural Comput, 7(6), 1129–1159.

Bell, M. A., & Cuevas, K. (2012). Using EEG to Study Cognitive Development: Issues and Practices. Journal of Cognition and Development, 13(3), 281–294. doi:10.1080/15248372.2012.691143

Bender, R., & Lange, S. (2001). Adjusting for multiple testing—when and how? Journal of Clinical Epidemiology, 54(4), 343–349. 10.1016/S0895-4356(00)00314-0

Berger, H. (1929). Über das Elektrenkephalogramm des Menschen. Archiv für Psychiatrie und Nervenkrankheiten, 87(1), 527–570. doi:10.1007/bf01797193

Bertrand, A. (2015). Distributed Signal Processing for Wireless EEG Sensor Networks. IEEE Transactions on Neural Systems and Rehabilitation Engineering, 23(6), 923–935. doi:10.1109/TNSRE.2015.2418351

Bigdely-Shamlo, N., Mullen, T., Kothe, C., Su, K. M., & Robbins, K. A. (2015). The PREP pipeline: standardized preprocessing for large-scale EEG analysis. Front Neuroinform, 9, 16. doi:10.3389/fninf.2015.00016

Borowicz, A. (2018). Using a multichannel Wiener filter to remove eye-blink artifacts from EEG data. Biomedical Signal Processing and Control, 45, 246–255. doi:10.1016/j.bspc.2018.05.012

Brooker, R. J., Bates, J. E., Buss, K. A., Canen, M. J., Dennis-Tiwary, T. A., Gatzke-Kopp, L. M., . . . Schmidt, L. A. (2020). Conducting Event-Related Potential (ERP) Research with Young Children: A Review of Components, Special Considerations and Recommendations for Research on Cognition and Emotion. J Psychophysiol, 34(3), 137–158. doi:10.1027/0269-8803/a000243

Buzzell, G. A., Morales, S., Valadez, E. A., Hunnius, S., & Fox, N. A. (2023). Maximizing the potential of EEG as a developmental neuroscience tool. Dev Cogn Neurosci, 60, 101201. doi:10.1016/j.dcn.2023.101201

Castellanos, N. P., & Makarov, V. A. (2006). Recovering EEG brain signals: artifact suppression with wavelet enhanced independent component analysis. J Neurosci Methods, 158(2), 300–312. doi:10.1016/j.jneumeth.2006.05.033

Cellier, D., Riddle, J., Petersen, I., & Hwang, K. (2021). The development of theta and alpha neural oscillations from ages 3 to 24 years. Dev Cogn Neurosci. doi:10.1016/j.dcn.2021.100969

Chang, C. Y., Hsu, S. H., Pion-Tonachini, L., & Jung, T. P. (2020). Evaluation of Artifact Subspace Reconstruction for Automatic Artifact Components Removal in Multi-Channel EEG Recordings. IEEE Transactions on Biomedical Engineering, 67(4), 1114–1121. doi:10.1109/TBME.2019.2930186

Daube, J. R. (2009). Waveforms and artifacts. In J. R. Daube & D. I. Rubin (Eds.), Clinical Neurophysiology (Third Edition ed., pp. 103–1144). USA: Oxford University Press.

De Blasio, F. M., & Barry, R. J. (2023). It’s time to RELAX and smell the roses! Clin Neurophysiol, 149, 176–177. doi:10.1016/j.clinph.2023.02.169

Debnath, R., Buzzell, G. A., Morales, S., Bowers, M. E., Leach, S. C., & Fox, N. A. (2020). The Maryland analysis of developmental EEG (MADE) pipeline. Psychophysiology, 57(6), e13580. doi:10.1111/psyp.13580

Delorme, A. (2023). EEG is better left alone. Sci Rep, 13(1), 2372. doi:10.1038/s41598-023-27528-0

Delorme, A., & Makeig, S. (2004). EEGLAB: an open source toolbox for analysis of single-trial EEG dynamics including independent component analysis. J Neurosci Methods, 134(1), 9-21. doi:10.1016/j.jneumeth.2003.10.009

Dimigen, O. (2020). Optimizing the ICA-based removal of ocular EEG artifacts from free viewing experiments. Neuroimage, 207, 116117. doi:10.1016/j.neuroimage.2019.116117

DiStefano, C., Dickinson, A., Baker, E., & Jeste, S. S. (2019). EEG Data Collection in Children with ASD: The Role of State in Data Quality and Spectral Power. Res Autism Spectr Disord, 57, 132–144. doi:10.1016/j.rasd.2018.10.001

Donoghue, T., Haller, M., Peterson, E. J., Varma, P., Sebastian, P., Gao, R., . . . Voytek, B. (2020). Parameterizing neural power spectra into periodic and aperiodic components. Nat Neurosci, 23(12), 1655–1665. doi:10.1038/s41593-020-00744-x

Edgar, J. C., Franzen, R. E., McNamee, M., Green, H. L., Shen, G., DiPiero, M., . . . Chen, Y. (2023). A comparison of resting-state eyes-closed and dark-room alpha-band activity in children. Psychophysiology, 60(6), e14285. doi:10.1111/psyp.14285

Fitzgibbon, S. P., DeLosAngeles, D., Lewis, T. W., Powers, D. M., Grummett, T. S., Whitham, E. M., . . . Pope, K. J. (2016). Automatic determination of EMG-contaminated components and validation of independent component analysis using EEG during pharmacologic paralysis. Clin Neurophysiol, 127(3), 1781–1793. doi:10.1016/j.clinph.2015.12.009

Fitzgibbon, S. P., Powers, D. M. W., Pope, K. J., & Clark, C. R. (2007). Removal of EEG Noise and Artifact Using Blind Source Separation. Journal of Clinical Neurophysiology, 24(3), 232–243. doi:10.1097/WNP.0b013e3180556926

Flo, A., Gennari, G., Benjamin, L., & Dehaene-Lambertz, G. (2022). Automated Pipeline for Infants Continuous EEG (APICE): A flexible pipeline for developmental cognitive studies. Dev Cogn Neurosci, 54, 101077. doi:10.1016/j.dcn.2022.101077

Frank, G., Makeig, S., & Delorme, A. (2022). A Framework to Evaluate Independent Component Analysis applied to EEG signal: testing on the Picard algorithm. arXiv. doi:arXiv:2210.08409

Gabard-Durnam, L. J., Mendez Leal, A. S., Wilkinson, C. L., & Levin, A. R. (2018). The Harvard Automated Processing Pipeline for Electroencephalography (HAPPE): Standardized Processing Software for Developmental and High-Artifact Data. Front Neurosci, 12, 97. doi:10.3389/fnins.2018.00097

Gabard-Durnam, L. J., Wilkinson, C., Kapur, K., Tager-Flusberg, H., Levin, A. R., & Nelson, C. A. (2019). Longitudinal EEG power in the first postnatal year differentiates autism outcomes. Nature Communications, 10(1), 4188. doi:10.1038/s41467-019-12202-9

Goncharova, I. I., & Barlow, J. S. (1990). Changes in EEG mean frequency and spectral purity during spontaneous alpha blocking. Electroencephalography and Clinical Neurophysiology, 76(3), 197–204. 10.1016/0013-4694(90)90015-C

Herve, E., Mento, G., Desnous, B., & Francois, C. (2022). Challenges and new perspectives of developmental cognitive EEG studies. Neuroimage, 260, 119508. doi:10.1016/j.neuroimage.2022.119508

Hochberg, Y. (1988). A sharper Bonferroni procedure for multiple tests of significance. Biometrika, 75(4), 800–802. doi:10.1093/biomet/75.4.800

Hrachovy, R. A., & Mizrahi, E. M. (2016). Atlas of Neonatal Electroencephalography (Fourth edition ed.). New York: Demos Medical.

Janani, A. S., Grummett, T. S., Bakhshayesh, H., Lewis, T. W., Willoughby, J. O., & Pope, K. J. (2018, 3-7 Sept. 2018). How Many Channels are Enough? Evaluation of Tonic Cranial Muscle Artefact Reduction Using ICA with Different Numbers of EEG Channels. Paper presented at the 2018 26th European Signal Processing Conference (EUSIPCO).

Jas, M., Engemann, D. A., Bekhti, Y., Raimondo, F., & Gramfort, A. (2017). Autoreject: Automated artifact rejection for MEG and EEG data. Neuroimage, 159, 417–429. doi:10.1016/j.neuroimage.2017.06.030

JASP Team. (2023). JASP (Version 0.18.1). [Computer software].

Jiang, X., Bian, G. B., & Tian, Z. (2019). Removal of Artifacts from EEG Signals: A Review. Sensors, 19(5). doi:10.3390/s19050987

Kirschfeld, K. (2005). The physical basis of alpha waves in the electroencephalogram and the origin of the “Berger effect”. Biological Cybernetics, 92(3), 177–185. doi:10.1007/s00422-005-0547-1

Klug, M., & Gramann, K. (2021). Identifying key factors for improving ICA-based decomposition of EEG data in mobile and stationary experiments. Eur J Neurosci, 54(12), 8406–8420. doi:10.1111/ejn.14992

Leach, S. C., Morales, S., Bowers, M. E., Buzzell, G. A., Debnath, R., Beall, D., & Fox, N. A. (2020). Adjusting ADJUST: Optimizing the ADJUST algorithm for pediatric data using geodesic nets. Psychophysiology, 57(8), e13566. doi:10.1111/psyp.13566

Lew, B. J., Fitzgerald, E. E., Ott, L. R., Penhale, S. H., & Wilson, T. W. (2021). Three-year reliability of MEG resting-state oscillatory power. Neuroimage, 243, 118516. doi:10.1016/j.neuroimage.2021.118516

Lopez, K. L., Monachino, A. D., Morales, S., Leach, S. C., Bowers, M. E., & Gabard-Durnam, L. J. (2022). HAPPILEE: HAPPE In Low Electrode Electroencephalography, a standardized pre-processing software for lower density recordings. Neuroimage, 260, 119390. doi:10.1016/j.neuroimage.2022.119390

Mair, P., & Wilcox, R. (2020). Robust statistical methods in R using the WRS2 package. Behav Res Methods, 52(2), 464–488. doi:10.3758/s13428-019-01246-w

Marshall, P. J., Bar-Haim, Y., & Fox, N. A. (2002). Development of the EEG from 5 months to 4 years of age. Clinical Neurophysiology, 113(8), 1199–1208. 10.1016/S1388-2457(02)00163-3

Mumtaz, W., Rasheed, S., & Irfan, A. (2021). Review of challenges associated with the EEG artifact removal methods. Biomedical Signal Processing and Control, 68. doi:10.1016/j.bspc.2021.102741

Niedermeyer, E. (1997). Alpha rhythms as physiological and abnormal phenomena. International Journal of Psychophysiology, 26(1), 31–49. 10.1016/S0167-8760(97)00754-X

R Core Team. (2020). R: A Language Environment for Statistical Computing. Vienna, Austria: R Foundation for Statistical Computing.

Robbins, K. A., Touryan, J., Mullen, T., Kothe, C., & Bigdely-Shamlo, N. (2020). How Sensitive Are EEG Results to Preprocessing Methods: A Benchmarking Study. IEEE Transactions on Neural Systems and Rehabilitation Engineering, 28(5), 1081–1090. doi:10.1109/TNSRE.2020.2980223

Roy, V., Shukla, P. K., Gupta, A. K., Goel, V., Shukla, P. K., & Shukla, S. (2021). Taxonomy on EEG Artifacts Removal Methods, Issues, and Healthcare Applications. Journal of Organizational and End User Computing (JOEUC), 33(1), 19–46. doi:10.4018/JOEUC.2021010102

Sazgar, M., & Young, M. G. (2019). EEG Artifacts. In Absolute Epilepsy and EEG Rotation Review. Switzerald Springer.

Somers, B., & Bertrand, A. (2016). Removal of eye blink artifacts in wireless EEG sensor networks using reduced-bandwidth canonical correlation analysis. J Neural Eng, 13(6), 066008. doi:10.1088/1741-2560/13/6/066008

Somers, B., Francart, T., & Bertrand, A. (2018). A generic EEG artifact removal algorithm based on the multi- channel Wiener filter. J Neural Eng, 15(3), 036007. doi:10.1088/1741-2552/aaac92

Tandle, A., & Jog, N. (2016). Classification of Artefacts in EEG Signal Recordings and EOG Artefact Removal using EOG Subtraction. Communications on Applied Electronics, 4, 12–19.

