## Supplementary Material for "RELAX-Jr: An Automated Pre-Processing Pipeline for Developmental EEG Recordings"

**Table S1:** Bayesian Paired-Sample T-Tests comparing alpha power (RMS values) between the eyes-open and eyes-closed recording conditions.

| Measure 1 | Measure 2 | $BF_{10}$ |
|---|---|---|
| MWF_wICA_EO | MWF_wICA_EC | $1.124\times10^{+34}$ |
| MWF_ONLY_EO | MWF_ONLY_EC | $9.521\times10^{+34}$ |
| wICA_ADJUST_EO | wICA_ADJUST_EC | $3.166\times10^{+37}$ |
| ICA_SUBTRACT_EO | ICA_SUBTRACT_EC | $1.911\times10^{+37}$ |
| MADE_EO | MADE_EC | $7.537\times10^{+37}$ |
| APICE_EO | APICE_EC | $2.043\times10^{+38}$ |
| HAPPE_EO | HAPPE_EC | $1.815\times10^{+16}$ |
| ASR_EO | ASR_EC | $1.404\times10^{+34}$ |

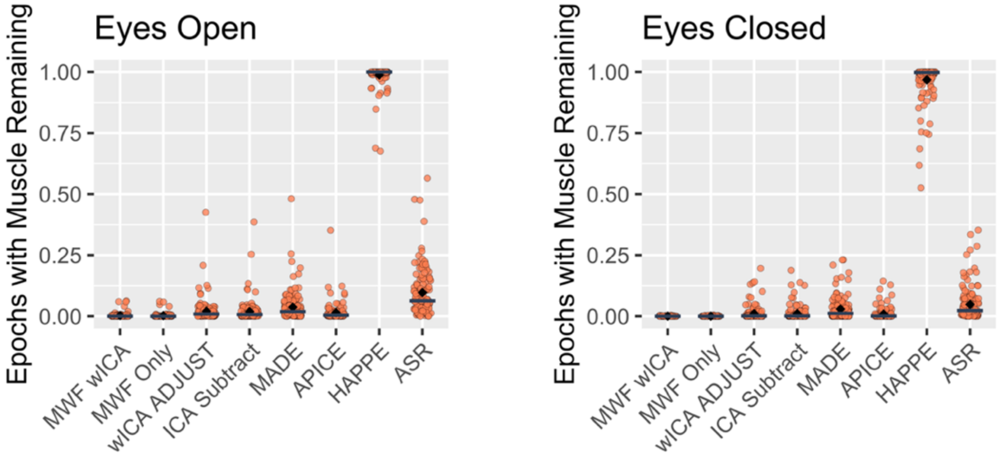


**Figure S1:** Proportion of epochs showing log-power log-frequency values above the -0.59 threshold for each of the pre-processing pipelines. Lower values reflect more effective cleaning of EMG activity. Un-winsorized values are shown (to compliment the Winsorized values presented in the manuscript). Results for the HAPPE pipeline are also included, which showed a high proportion of epochs containing EMG compared to the other pipelines.

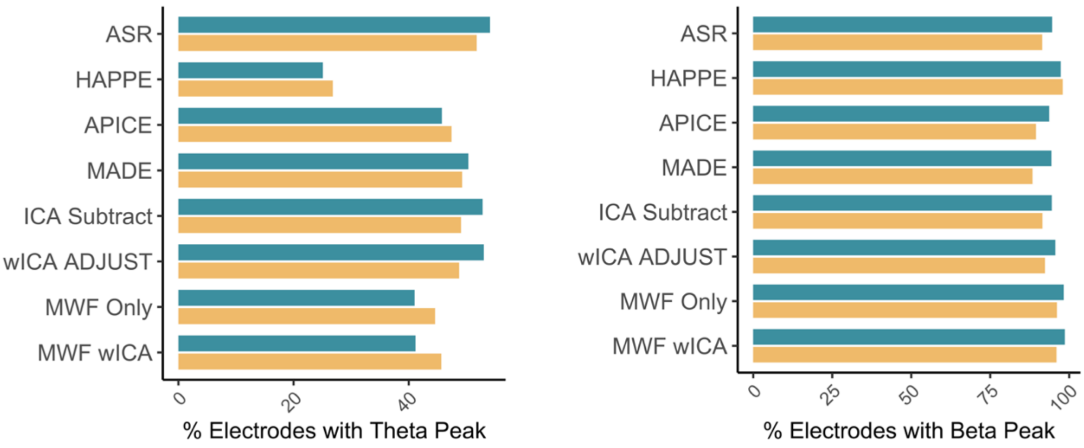


**Figure S2:** Percentage of total electrodes having a detected peak within the theta (4-7 Hz; left) and beta (13-30 Hz; right) bands after removal of the 1/*f*-like aperiodic signal for the eyes-open and eyes-closed data after cleaning with each pipeline (average over all participants). Given the strong performance of ASR, wICA Adjust, and ICA Subtract for theta peak detection across electrodes, these specific pipelines might be particularly useful if theta oscillations are the focus for analysis.

**Cleaned Dataset Examples**

The below figures provide examples from a single participant (eyes-open recording) of the data cleaned using the various pipelines. On the left is an example 5-second data segment and on the right is the spectral plot from EEGLAB. Note, all data are continuous files taken prior to segmentation or interpolation of any bad channels that might have been removed.

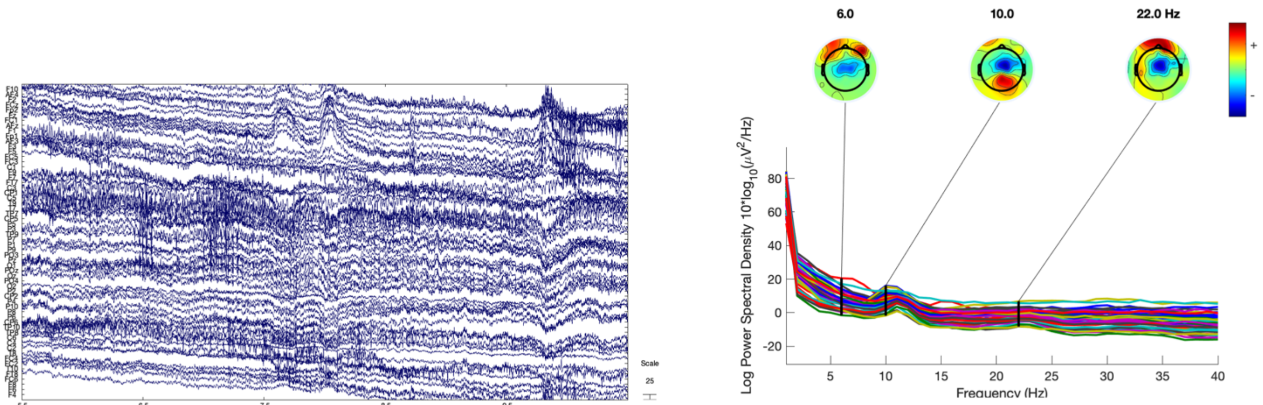


**Figure S3:** Raw EEG. No cleaning applied. DC offset removed to aid visualisation.

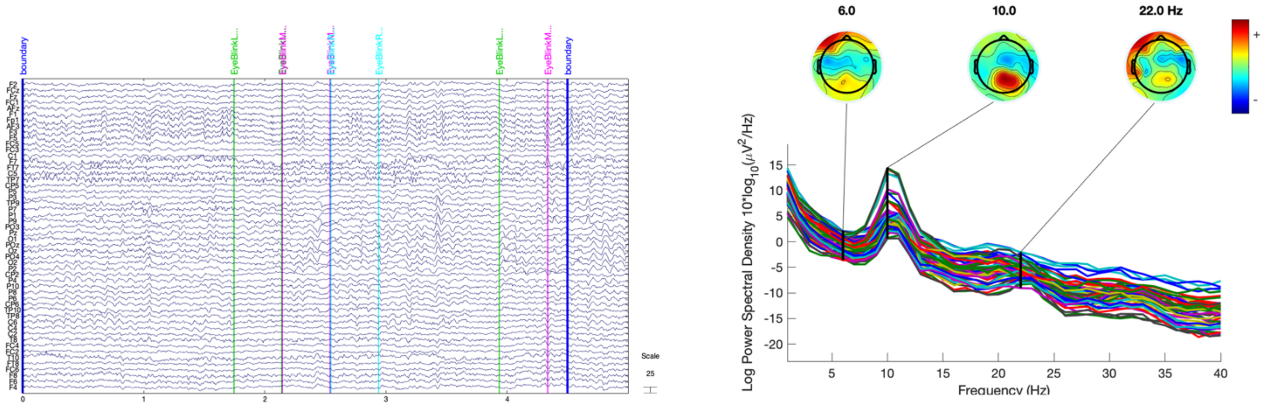


**Figure S4:** MWF wICA

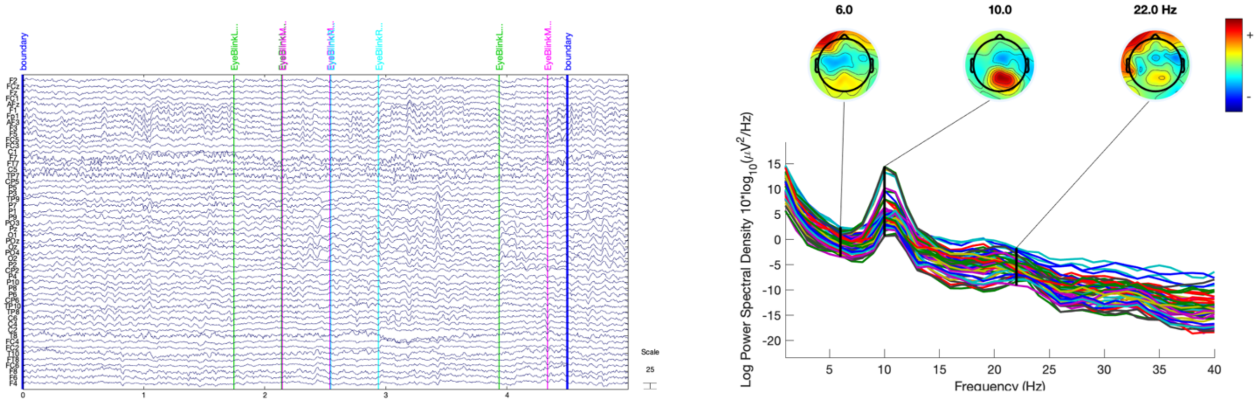


**Figure S5:** MWF Only

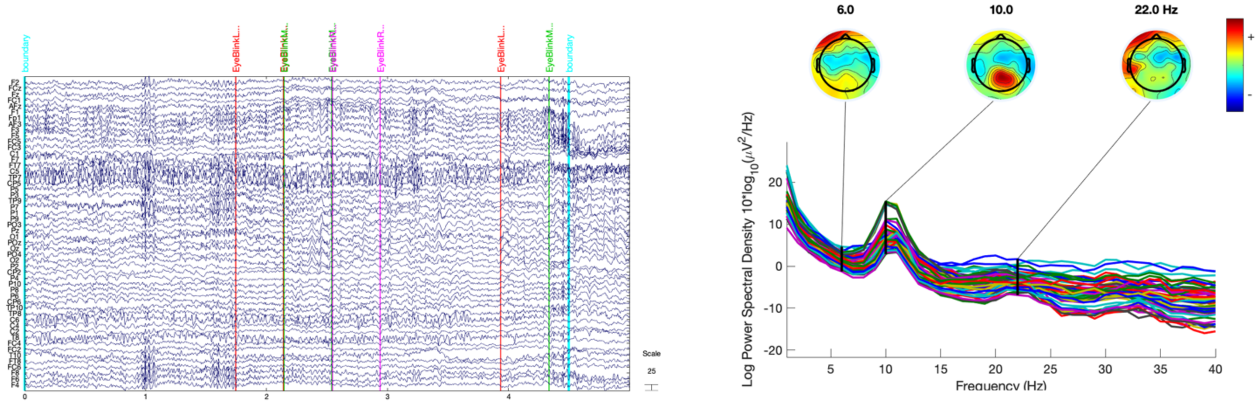


**Figure S6:** wICA ADJUST

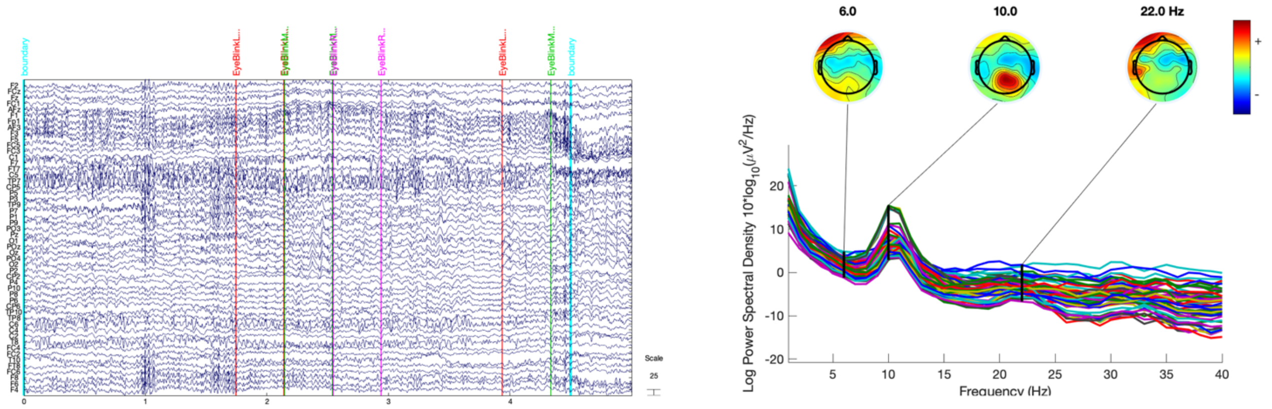


**Figure S7:** ICA Subtract

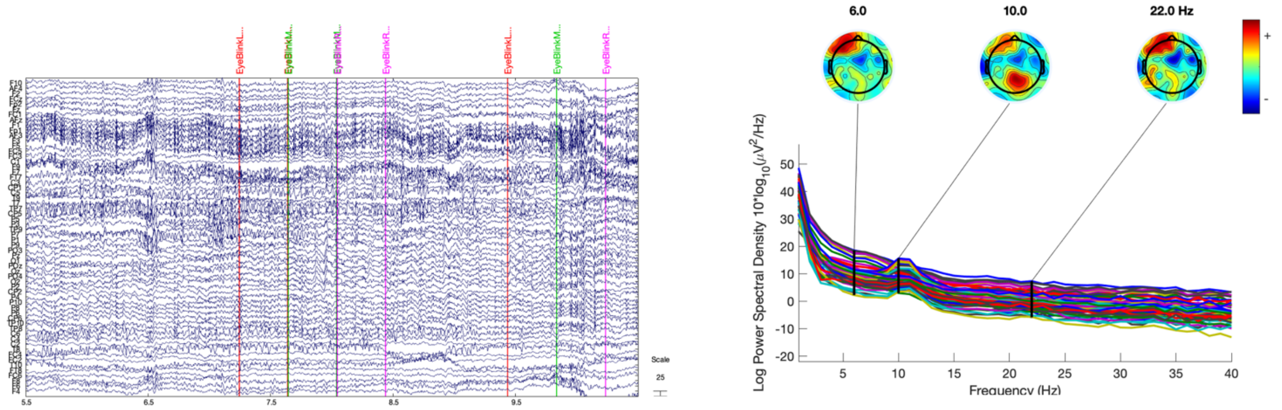


**Figure S8:** MADE

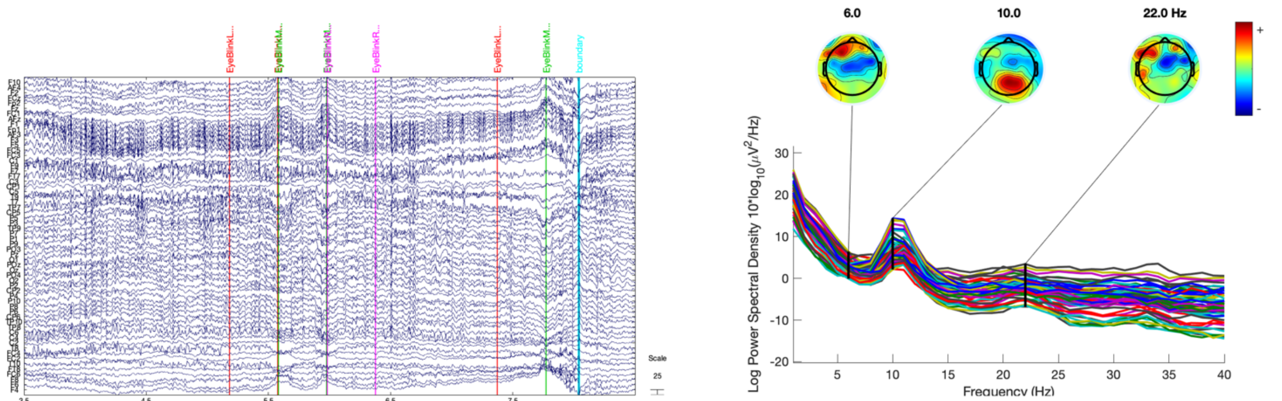


**Figure S9:** APICE

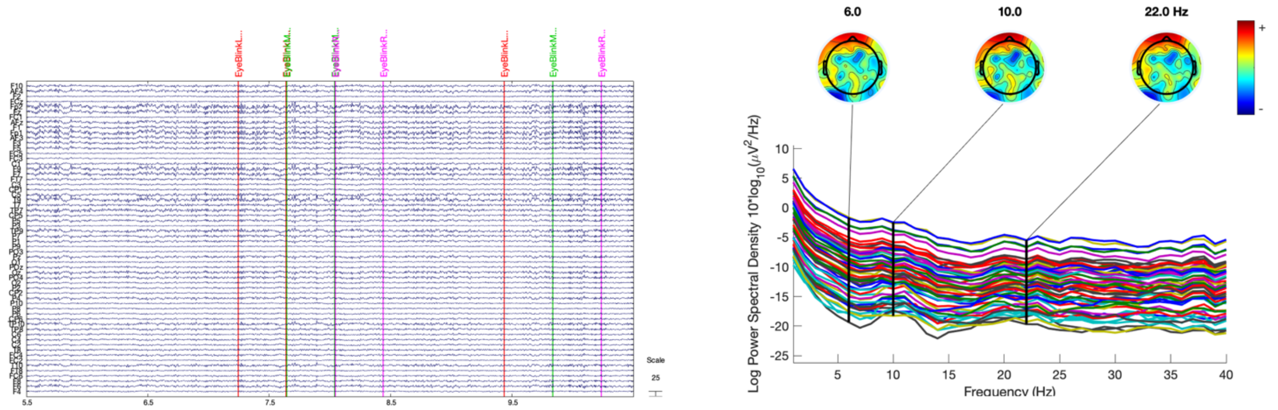


**Figure S10:** HAPPE

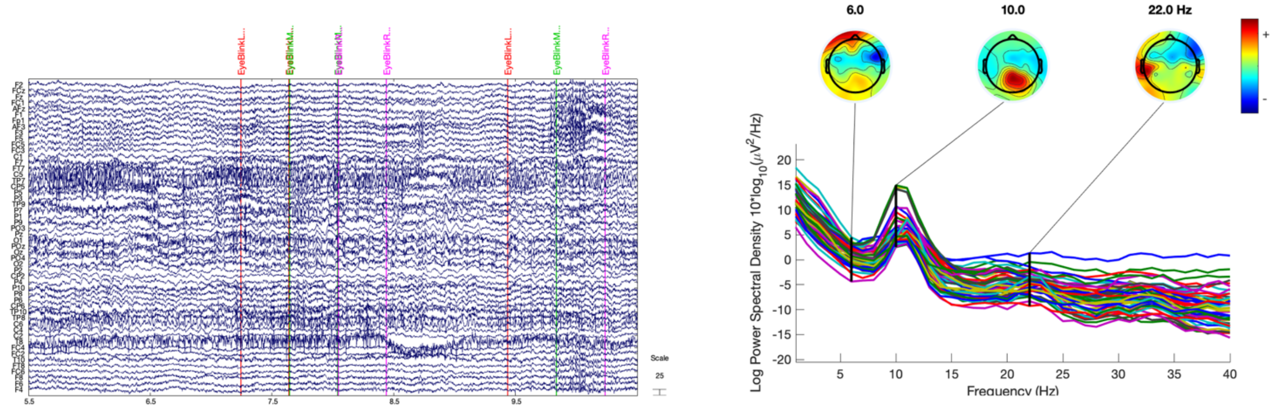


**Figure S11:** ASR

### Post-Hoc Tests

Figures below show the 95% confidence intervals for post-hoc comparisons run following the omnibus ANOVAs (reported in the manuscript), between each pipeline on the left (vertical) axis, and each pipeline along the bottom (horizontal) axis. Significant results after multiple comparison corrections (Hochberg) are denoted by an asterisk (*).

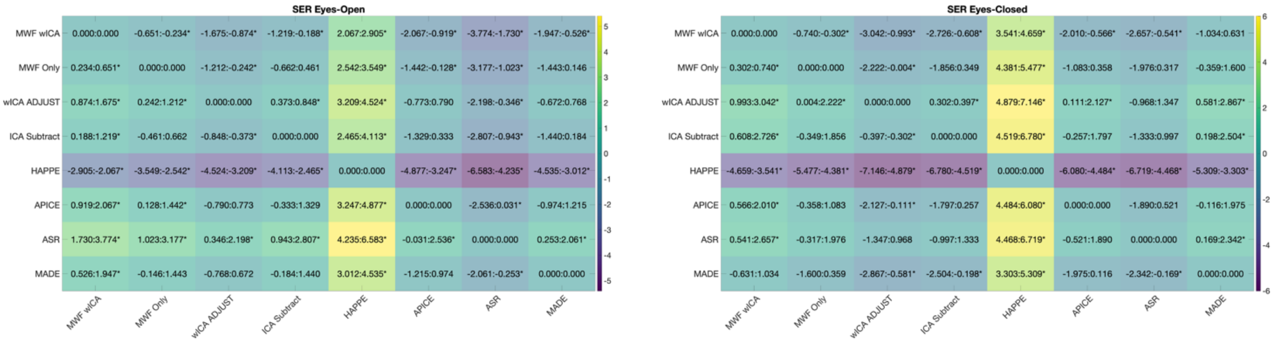

SER Eyes-Open

| | MWF wICA | MWF Only | wICA ADJUST | ICA Subtract | HAPPE | APICE | ASR | MADE |
|---|---|---|---|---|---|---|---|---|
| MWF wICA | 0.000:0.000 | -0.651:-0.234* | -1.675:-0.874* | -1.219:-0.188* | 2.067:2.905* | -2.067:-0.919* | -3.774:-1.730* | -1.947:-0.526* |
| MWF Only | 0.234:0.651* | 0.000:0.000 | -1.212:-0.242* | -0.662:0.461 | 2.542:3.549* | -1.442:-0.128* | -3.177:-1.023* | -1.443:0.146 |
| wICA ADJUST | 0.874:1.675* | 0.242:1.212* | 0.000:0.000 | 0.373:0.848* | 3.209:4.524* | -0.773:0.790 | -2.198:-0.346* | -0.672:0.768 |
| ICA Subtract | 0.188:1.219* | -0.461:0.662 | -0.848:-0.373* | 0.000:0.000 | 2.465:4.113* | -1.329:0.333 | -2.807:-0.943* | -1.440:0.184 |
| HAPPE | -2.905:-2.067* | -3.549:-2.542* | -4.524:-3.209* | -4.113:-2.465* | 0.000:0.000 | -4.877:-3.247* | -6.583:-4.235* | -4.535:-3.012* |
| APICE | 0.919:2.067* | 0.128:1.442* | -0.790:0.773 | -0.333:1.329 | 3.247:4.877* | 0.000:0.000 | -2.536:0.031* | -0.974:1.215 |
| ASR | 1.730:3.774* | 1.023:3.177* | 0.346:2.198* | 0.943:2.807* | 4.235:6.583* | -0.031:2.536* | 0.000:0.000 | 0.253:2.061* |
| MADE | 0.526:1.947* | -0.146:1.443 | -0.768:0.672 | -0.184:1.440 | 3.012:4.535* | -1.215:0.974 | -2.061:-0.253* | 0.000:0.000 |

SER Eyes-Closed

| | MWF wICA | MWF Only | wICA ADJUST | ICA Subtract | HAPPE | APICE | ASR | MADE |
|---|---|---|---|---|---|---|---|---|
| MWF wICA | 0.000:0.000 | -0.740:-0.302* | -3.042:-0.993* | -2.726:-0.608* | 3.541:4.659* | -2.010:-0.566* | -2.657:-0.541* | -1.034:0.631 |
| MWF Only | 0.302:0.740* | 0.000:0.000 | -2.222:-0.004* | -1.856:0.349 | 4.381:5.477* | -1.083:0.358 | -1.976:0.317 | -0.359:1.600 |
| wICA ADJUST | 0.993:3.042* | 0.004:2.222* | 0.000:0.000 | 0.302:0.397* | 4.879:7.146* | 0.111:2.127* | -0.968:1.347 | 0.581:2.867* |
| ICA Subtract | 0.608:2.726* | -0.349:1.856 | -0.397:-0.302* | 0.000:0.000 | 4.519:6.780* | -0.257:1.797 | -1.333:0.997 | 0.198:2.504* |
| HAPPE | -4.659:-3.541* | -5.477:-4.381* | -7.146:-4.879* | -6.780:-4.519* | 0.000:0.000 | -6.080:-4.484* | -6.719:-4.468* | -5.309:-3.303* |
| APICE | 0.566:2.010* | -0.358:1.083 | -2.127:-0.111* | -1.797:0.257 | 4.484:6.080* | 0.000:0.000 | -1.890:0.521 | -0.116:1.975 |
| ASR | 0.541:2.657* | -0.317:1.976 | -1.347:0.968 | -0.997:1.333 | 4.468:6.719* | -0.521:1.890 | 0.000:0.000 | 0.169:2.342* |
| MADE | -0.631:1.034 | -1.600:0.359 | -2.867:-0.581* | -2.504:-0.198* | 3.303:5.309* | -1.975:0.116 | -2.342:-0.169* | 0.000:0.000 |

**Figure S12:** SER eyes-open (left) and eyes-closed (right).

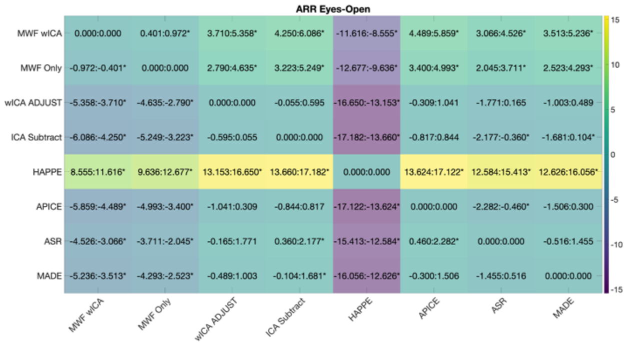

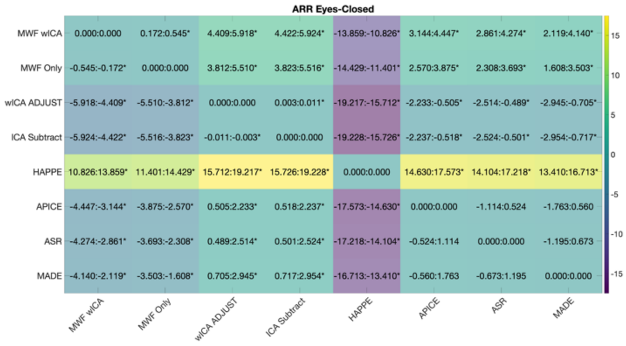


**Figure S13:** ARR eyes-open (left) and eyes-closed (right).

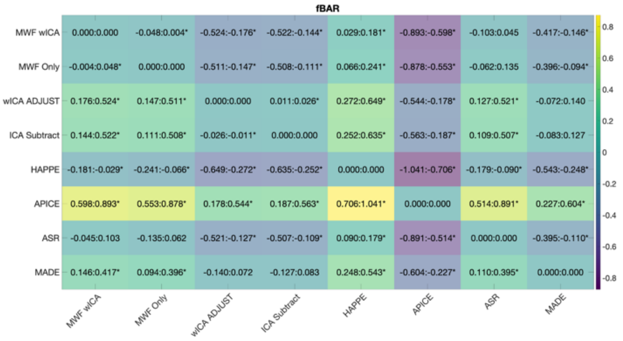

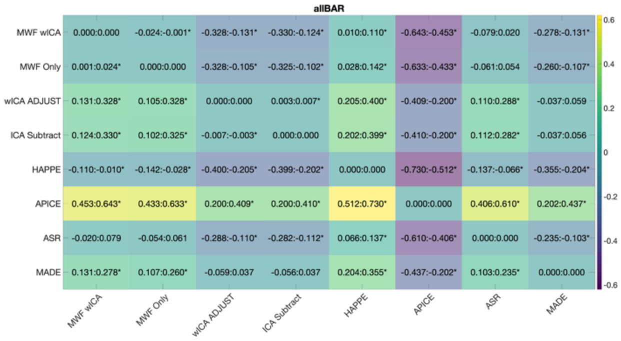


**Figure S14:** fBAR (left) and allBar (right) for the eyes-open data.

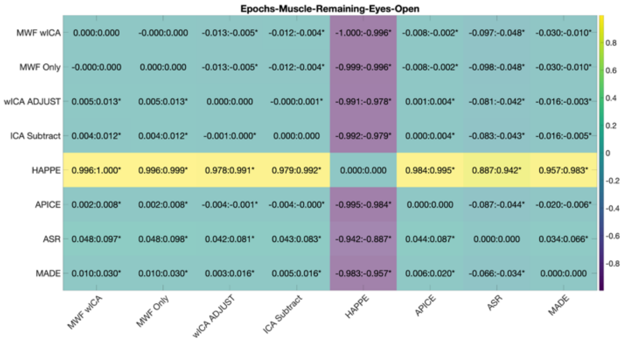

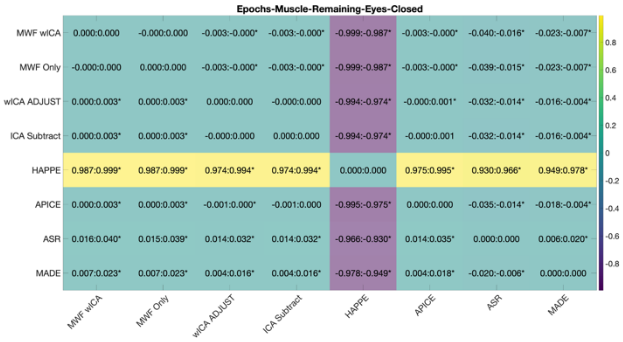


**Figure S15:** Epochs with muscle remaining after cleaning eyes-open (left) and eyes-closed (right).

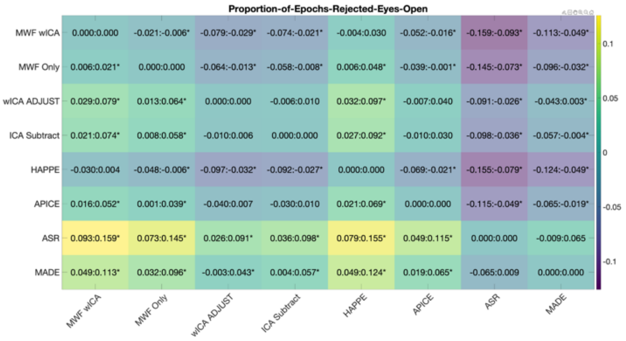

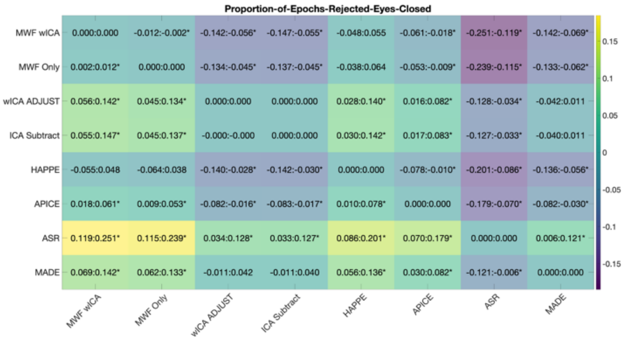


**Figure S16:** Proportion of epochs rejected during cleaning eyes-open (left) and eyes-closed (right).

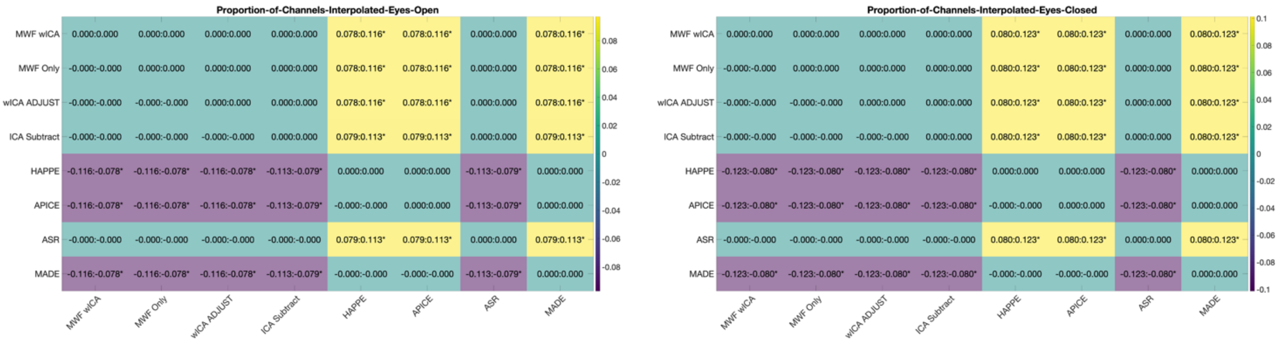

**Figure S17:** Proportion of channels interpolated eyes-open (left) and eyes-closed (right).
